# Self-propagating organelle biogenesis on synthetic beads

**DOI:** 10.64898/2026.01.09.698636

**Authors:** Min Peng, Siu-Shing Wong, Bocheng Xiao, Lorena Blaya-Martinez, Liuyi Hu, Zhe Feng, Nada Mohamad, Mehadi Hasan, Saroj Saurya, Zsofi Novak, Joao M. Monteiro, Tess Harrison, Chia-Chun Chang, Jordan W. Raff

## Abstract

Graphical Abstract

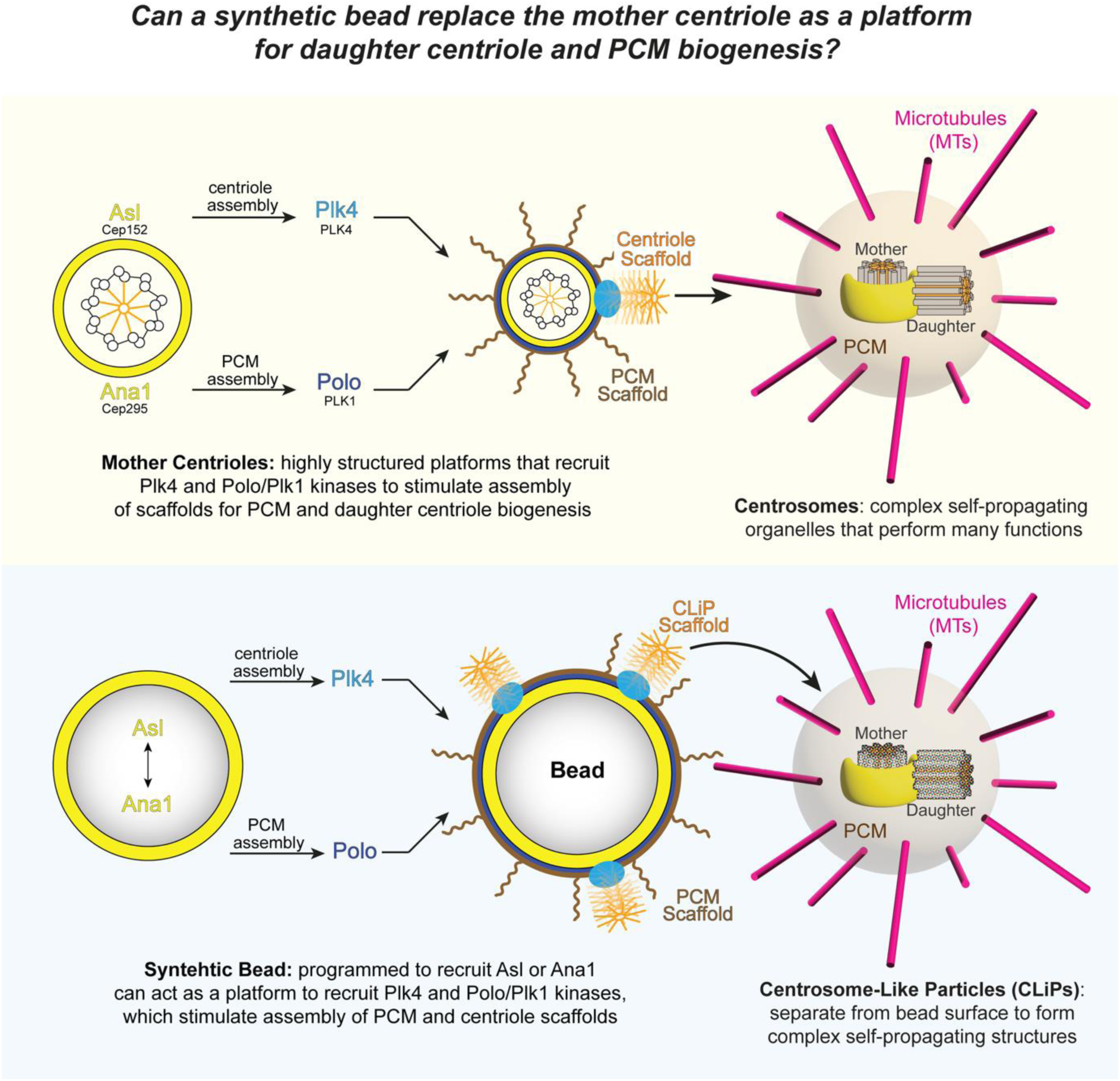

How cells build their many organelles remains poorly understood. Centrosomes are a striking example: structurally complex organelles built from more than one hundred different proteins. Here we show that a synthetic bead can generate centrosome-like particles (CLiPs) in early *Drosophila* embryos if it is programmed to recruit any one of five essential centrosome assembly proteins. CLiPs behave remarkably like centrosomes: organising centrosomal proteins, microtubules and actomyosin, forming spindle poles, and undergoing multiple rounds of duplication—all in synchrony with the embryonic cell cycle. The beads generate CLiPs because they recruit Plk4, the master regulator of centrosome biogenesis, in a specific manner that stimulates its assembly activity. Thus, recruiting a single protein to a synthetic surface is sufficient to seed the self-assembly of a complex, self-propagating entity. These findings suggest a general framework for organelle biogenesis in which simple localised molecular inputs can seed the self-assembly of highly complex cellular structures.

## Introduction

Cells contain many organelles that perform a myriad of functions. How cells assemble such complex structures and regulate their biogenesis to maintain cell and organelle homeostasis is poorly understood ^1–3^. Centrosomes are an attractive model for studying organelle biogenesis as most proliferating animal cells inherit a single centrosome whose number, shape and size are tightly controlled ^4–6^. Centrosomes comprise a highly structured 9-fold symmetric pair of mother-daughter centrioles surrounded by a more amorphous matrix of Pericentriolar Material (PCM) (Figure 1A_i_) ^7^. In proliferating cells, centrosomes duplicate precisely once per cell cycle through a canonical pathway (Figure 1A) ^8,9^. Mother and daughter centrioles disengage (Figure 1A_ii_), the daughter is converted into a mother, and each mother assembles a single daughter that emerges orthogonally from its side during S-phase (Figure 1A_iii_). As cells enter mitosis, PCM expands around the mother centrioles (Figure 1A_iii_), enabling the duplicated centrosomes to form the poles of the mitotic spindle (Figure 1A_iv_).

**Figure 1.**
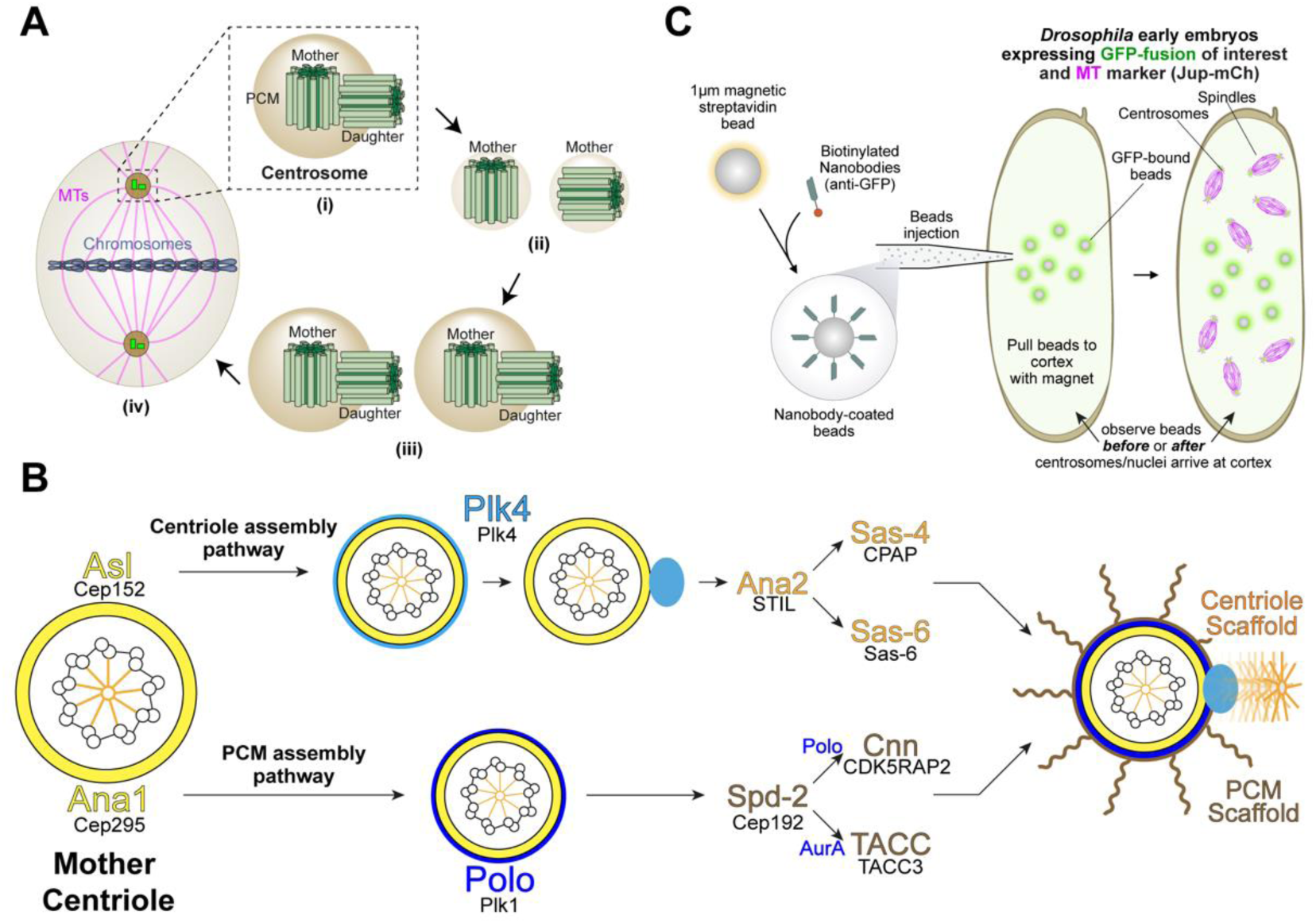
Centrosome assembly. (**A**) Illustration of centrosomes and canonical centrosome duplication. (**B**) The proposed pathways of daughter centriole and mitotic PCM assembly in *Drosophila* embryos. The names of human homologues are indicated below each protein. (**C**) Illustration of the synthetic bead injection assay.

Centrosomes contain multiple copies of more than 100 different types of protein ^10^, and how such complicated structures duplicate remains one of the central unresolved questions in cell biology. Unlike DNA replication, in which the information required to assemble a new strand is encoded within the parental double helix, it is unclear where the information that guides daughter centriole assembly resides. Crucially, centrioles can assemble *de novo* in certain species and specialised cell types, demonstrating that a pre-existing mother centriole is not an essential pre-requisite for centriole biogenesis ^11–16^. If, however, the *de novo* pathway is induced in proliferating cells by the removal of the pre-existing centrioles, this pathway lacks the precision of the canonical pathway, generating a variable number of centrioles over an extended and more variable time frame ^17–21^. Moreover, in some cases, the *de novo* pathway also generates centrioles with structural defects ^17,22^. Thus, the prevailing view is that mother centrioles are required to promote the relatively rapid, high-fidelity canonical duplication observed in proliferating cells ^23^. Whether they do this by providing a structural template is unclear, but it has been proposed that mother centrioles can “shed” their 9-fold symmetric inner cartwheel structure, which then binds to the side of the mother to template canonical duplication ^24,25^.

Despite their complexity, only a surprisingly small number of conserved proteins are essential for mother centrioles to generate daughters and to assemble an expanded PCM during mitosis ^26–28^. In *Drosophila* embryos, the centriole and PCM assembly pathways share a similar logic (Figure 1B). Asl/Cep152 and Ana1/Cep295 (fly/human nomenclature, used throughout) each form a torus around the mother centriole (*yellow*, Figure 1B). Each then recruits a protein kinase that functions as the master regulator of centriole (Plk4/PLK4) or mitotic PCM (Polo/PLK1) biogenesis ^29,30^ (*light/dark blue*, Figure 1B). For centriole biogenesis, Plk4 breaks symmetry to establish a single site on the mother from where it phosphorylates Ana2/STIL to promote interactions with Sas-6/SAS-6 and Sas-4/CPAP—leading to the assembly of the ninefold symmetric inner cartwheel scaffold that supports centriole assembly ^31–36^ (*orange*, Figure 1B). For PCM biogenesis, Polo phosphorylates Spd-2/Cep192 to promote interactions with Cnn/CDK5RAP2 and Polo or TACC/TACC3 and Aurora A/AURKA to generate a scaffold that extends around the mother to support mitotic PCM assembly ^37,38^ (*brown*, Figure 1B).

In this framework, mother centrioles may not need to provide a structural template but rather may simply function as platforms that support the self-assembly of simple scaffolds that guide canonical assembly. If so, we reasoned it should be possible to reconstitute canonical centrosome biogenesis on an entirely synthetic platform.

Here we test this idea by programming synthetic beads to recruit individual centrosome assembly proteins following injection into early *Drosophila* embryos, where centrosomes normally undergo 13 rounds of rapid, near-synchronous canonical duplication within a common syncytial cytoplasm^39^. We find that beads programmed to recruit Plk4 or Polo/Plk1 did not generate centrioles or recruit PCM, but beads programmed to recruit any one of Asl/Cep152, Ana1/Cep295, Ana2/STIL, Sas-6, or Sas-4/CPAP generated PCM and also **C**entrosome-**Li**ke **P**article**s** (**CLiPs**), some of which appear to be functionally indistinguishable from the endogenous centrosomes. We show that these beads generate CLiPs and PCM because they recruit Asl/Cep152 and Ana1/Cep295, which then recruit and activate Plk4 and Polo to initiate CLiP and PCM biogenesis, respectively. Thus, a synthetic bead can support the assembly of a complex self-propagating structure simply by recruiting a single biogenesis-promoting protein.

## Results

### A synthetic platform to support centrosome biogenesis

We wanted to test whether a synthetic bead programmed to recruit core assembly proteins could replace the mother centriole as a platform for centrosome biogenesis in the early *Drosophila* embryo. To do this, we coupled biotinylated anti-GFP Nanobodies—or, in some experiments, anti-ALFA-tag Nanobodies—to 1μm streptavidin magnetic beads *in vitro*. We then injected these beads into early embryos that expressed various individual GFP fusions (or ALFA-tagged fusions) that bind to the Nanobody beads, as well as Jupiter-mCherry (Jup-mCh) as a microtubule (MT) marker (Figure 1C). We used a magnet to position the beads at the embryo cortex so we could compare their behaviour to the endogenous centrosomes that arrived at the embryo cortex during nuclear cycle 10 (NC10) (Figure 1C).

### Synthetic beads coupled to Sas-6 and Ana2, but not Plk4 or Polo/PLK1, can organise PCM and MTs

We first asked whether recruiting either of the master regulators of centrosome biogenesis, Plk4 or Polo, was sufficient to initiate centriole and/or PCM assembly. GFP-Nanobody beads injected into embryos expressing either GFP-Plk4 or Polo-GFP efficiently recruited each protein to their surface but failed to organise MT arrays or generate centrosomes (Figure 2A,B; Video S1, S2).

**Figure 2.**
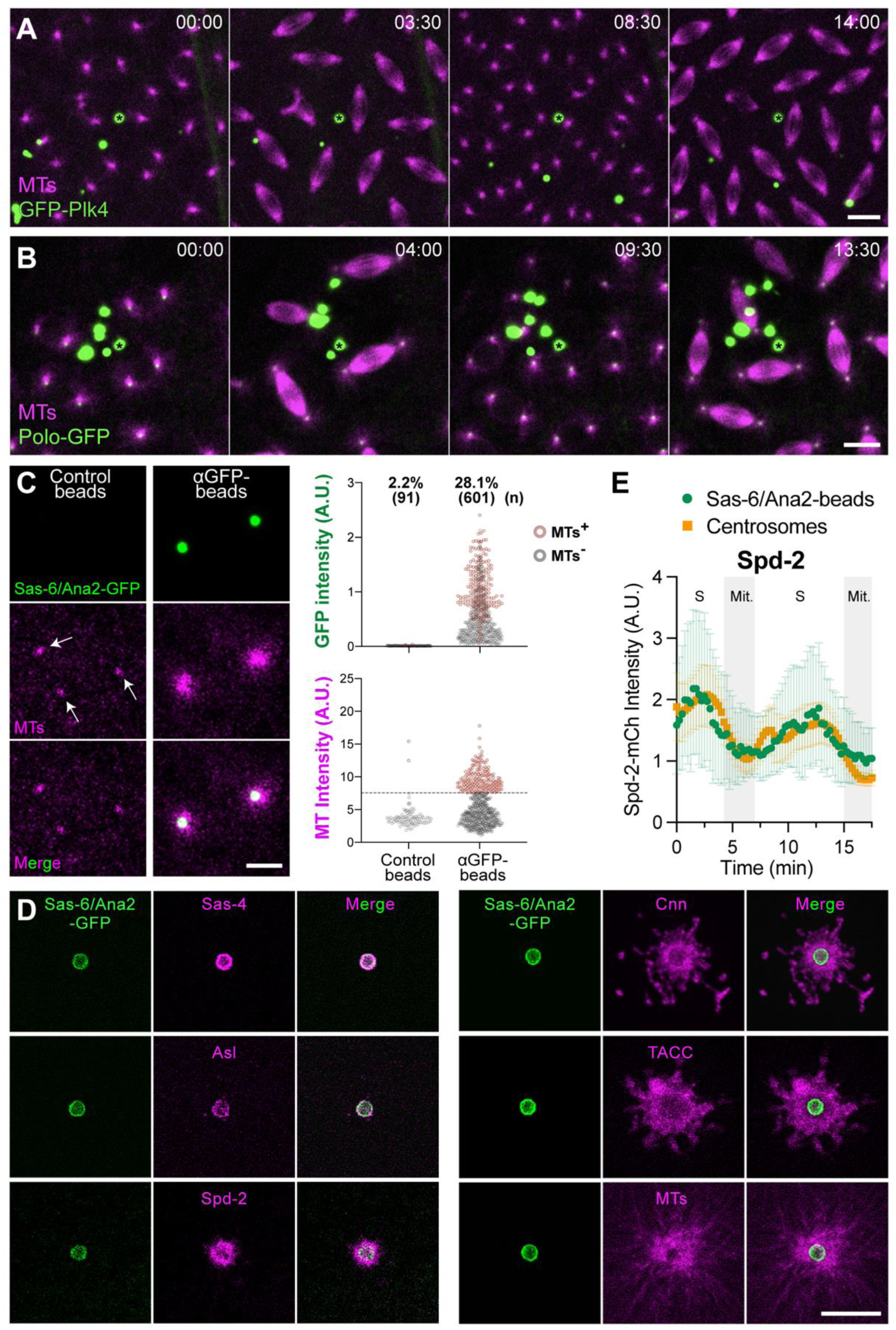
Plk4 and Polo beads cannot organise MTs; Sas-6/Ana2 beads can. (**A,B**) Images show stills from videos of embryos expressing GFP-Plk4 (A) or Polo-GFP (B) (*green*) and Jup-mCh (MTs) (*magenta*) injected with anti-GFP-Nanobody beads. Time (mins:secs) is indicated relative to the first image (t=00:00). These beads recruit large amounts of each GFP-fusion, but they do not detectably organise MTs or generate centrosomes (N=4-18 embryos; n≥200 beads analysed; see Figure S7 for quantification). (**C**) Images show anti-ALFA-Nanobody (Control) or anti-GFP-Nanobody coupled beads ∼20-30mins after injection into an embryo expressing Sas-6-GFP, Ana2-GFP (*green*) and Jup-mCh (MTs) (*magenta*). The GFP beads recruit the GFP-fusions and organise MT arrays; control beads exhibit only weak autofluorescence in the Jup-mCh channel (*arrows*). Graphs quantify the intensity of GFP (top) or MTs (bottom) on individual beads. Beads above or below a threshold MT-fluorescence were scored as MT +ve (*brown dots*) or -ve (*grey dots*), respectively (see Figure S1 for methodology). The percentage of beads organising MTs (%) and the number of beads analysed (n) is indicated; N=5-15 embryos per condition. (**D**) Images show GFP-Nanobody beads ∼20-30min after injection into embryos expressing Sas-6-GFP, Ana2-GFP (*green*) and red-fluorescent fusions to various individual centriole/centrosome proteins, as indicated (*magenta*). **(E)** Graph quantifies the fluorescence intensity (mean±SD) of the PCM-scaffold protein Spd-2-mCh at centrosomes (*orange squares*) and at Sas-6/Ana2 beads (*green dots*) as a typical embryo proceeded through NC11-12; (Mit) indicates mitosis; (S) indicates S-phase. Scale bars = 10µm (A,B); 5µm (C,D).

We next tested whether the simultaneous recruitment of Sas-6 and Ana2 could promote centrosome assembly, as we had previously shown that when co-overexpressed in embryos these proteins form **S**as-6/**A**na2 **P**article**s** (**SAPs**) that recruit PCM and organise MTs ^40^. GFP-Nanobody beads injected into early embryos expressing Sas-6-GFP and Ana2-GFP (at endogenous levels, so no SAPs were formed) could organise extensive MT arrays (Figure 2C) (see Figure S1 for an explanation of the computational methods used for quantification).

To determine whether this MT organisation reflected PCM recruitment, we generated Sas-6/Ana2-GFP beads in embryos expressing red-fluorescent fusions of other individual centrosome proteins. The beads recruited each protein in a pattern that closely resembled its normal distribution at centrosomes ^40^ (Figure 2D). Importantly, PCM recruitment was not static. During each nuclear cycle in *Drosophila* embryos, centrioles generate a pulse of Polo/PLK1 and Spd-2 recruitment that drives mitotic-PCM expansion ^41^. Sas-6/Ana2 beads exhibited a similar oscillatory behaviour, with pulses of PCM recruitment occurring in phase with the endogenous centrosomes (Figure 2E). Thus, recruitment of Sas-6 and Ana2 is sufficient to endow synthetic beads with dynamic centrosome-like behaviour, allowing them to recruit PCM and organise MTs in a manner that closely mirrors endogenous centrosomes.

### Sas-6/Ana2 beads generate <u>C</u>entrosome-<u>Li</u>ke <u>P</u>articles (CLiPs)

Sas-6/Ana2 beads organised PCM and MTs within ∼20 min of injection into early embryos (t=0:00, Figure 3A; Video S3). As endogenous centrosomes began arriving at the cortex (*arrowhead*, t=17:30, Figure 3A), however, the beads also started to generate small particles that detached from their surface and organised their own MT arrays (*cyan asterisk*, t=17:30, Figure 3A). Remarkably, many of these particles proceeded to duplicate in synchrony with the endogenous centrosomes (Figure 3B; Video S4), often undergoing four successive rounds of duplication, before all duplication ceased as the embryo stopped cycling and began to cellularise (Figure 3C; Video S5) (see also Figure S2; Video S3).

**Figure 3.**
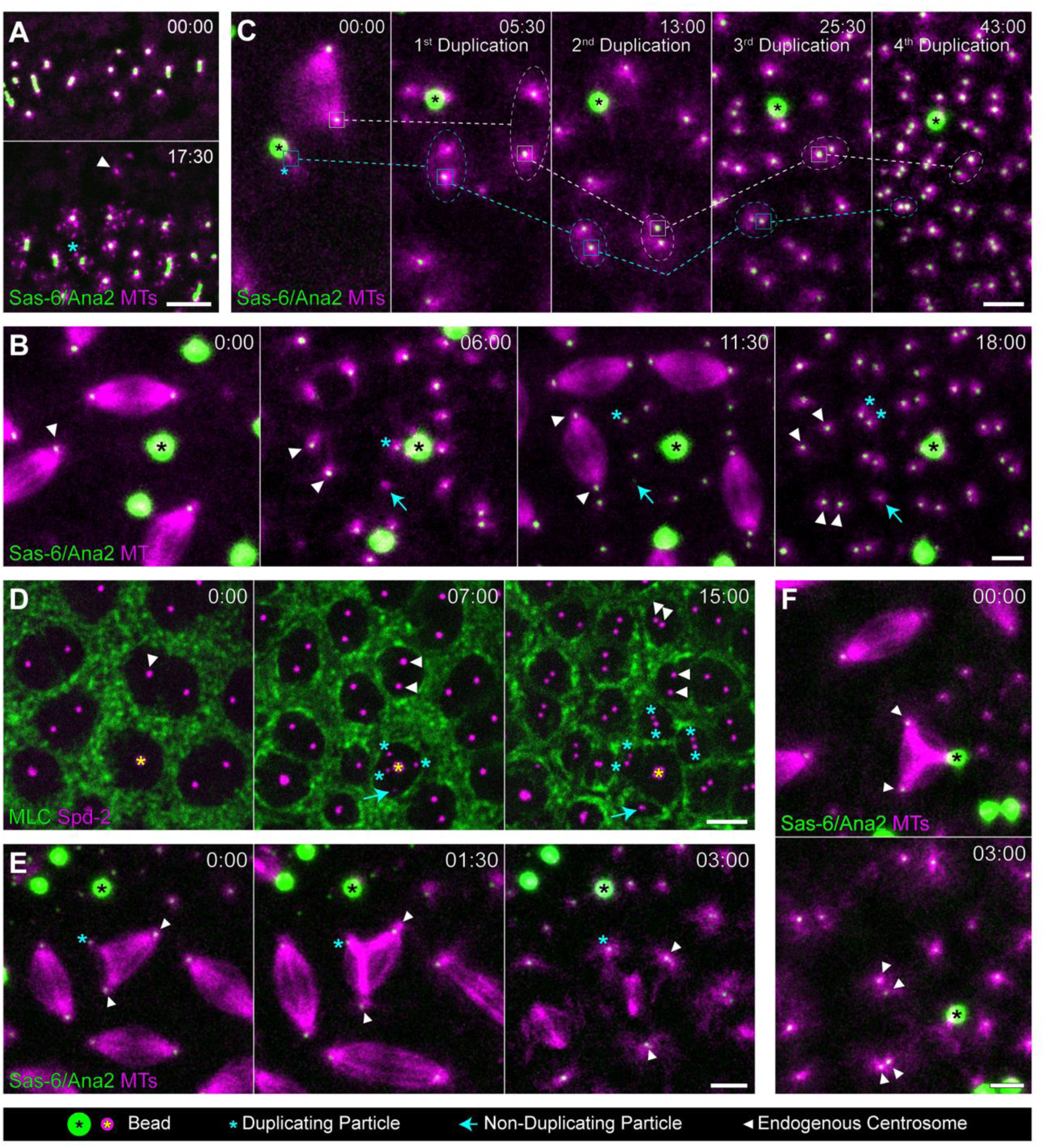
Sas-6/Ana2 beads generate <u>C</u>entrosome-<u>Li</u>ke <u>P</u>articles (CLiPs) that are functionally similar to centrosomes. **(A)** Image shows Sas-6/Ana2-GFP beads (*green*) organising MTs (*magenta*) prior to the arrival of the endogenous centrosomes at the embryo cortex (t=00:00). As the centrosomes reach the cortex (*arrowhead*, t=17:30) the beads generate MT-particles (*cyan asterisk*, t=17:30). **(B)** Images show Sas-6/Ana2-GFP beads (*black asterisk)* and endogenous spindles (*arrowhead* highlights an endogenous centrosome) (t=00:00). After mitosis the duplicated centrosomes separate, and several particles separate from the bead surface (t=6:00)—a bright (*cyan asterisk*) and a dim (*cyan arrow*) particle are highlighted. The embryo proceeds through mitosis (t=11:30), after which, the duplicated centrosomes and brighter particles separate, but the dimmer particle does not (t=18:00). **(C)** A Sas-6/Ana2-GFP bead generates a single particle that proceeds through four rounds of duplication in synchrony with the endogenous centrosomes. Solid boxes highlight a centrosome (*white*) and particle (*cyan*) that duplicated during each nuclear cycle—dashed ovals indicate the duplicated pair. (**D**) A Sas-6/Ana2-ALFA bead (*yellow asterisk*) organises a cortical actomyosin “cage” around itself—visualised here with Myosin Light Chain (MLC)-GFP (*green*) (t=00:00). The ALFA-tag is non-fluorescent, so the bead and centrosomes are visualised with Spd-2-mCherry (*magenta*). The bead generates three particles (t=07:00) that duplicate and organise their own actomyosin cages (t=15:00), and a dimmer particle (*cyan arrow*) that does not duplicate but still organises an actomyosin cage. **(E,F)** Images show a particle (E) or bead (F) that contacts a nearby spindle and forms a spindle pole that, at this resolution, appears functionally indistinguishable from the centrosome-organised spindle poles. Scale bars = 20 μm (A); 5 μm (B,C,E,F); 10 μm (D).

Not all these particles were duplication competent. Some initially contained low levels of Sas-6-GFP/Ana2-GFP and, although they organised MTs, such particles often failed to duplicate (e.g. *cyan arrows*, Figure 3B,D). Some of these particles eventually acquired more Sas-6-GFP and duplicated in later cycles, while others dissipated and/or were lost from view (Figure S3; Video S6).

As we describe below, many of these particles are very similar to centrosomes, so we collectively refer to them as **centrosome-like particles** (**CLiPs**). In the following experiments we often quantify the ability of various beads to organise MTs and use this as a proxy for CLiP generation, as the ability of beads to organise MTs was highly correlated with their ability to generate CLiPs. Approximately 80% of MT-organising Sas-6/Ana2 beads generated at least one, and usually multiple, CLiP(s), and we never observed a Sas-6/Ana2 bead that failed to organise MTs generate a CLiP (179 MT-generating beads analysed from nine embryos).

### CLiPs are functionally similar to centrosomes

Although most CLiPs did not form spindles, we reasoned that this might be because, like the endogenous centrosomes, CLiPs might organise a cortical actomyosin network around themselves that normally helps keep neighbouring centrosomes separated from one another in the syncytial embryo ^42,43^. To test this, we generated Sas-6/Ana2-ALFA beads in embryos expressing Spd-2-mCherry (Spd-2-mCh) (as a centrosome marker) and myosin light chain (MLC)-GFP. The CLiPs, and the Sas-6/Ana2 beads that generated them, behaved similarly to centrosomes, organising a cortical myosin network around themselves that prevented their MT arrays from interacting with each other and with the MTs associated with nearby centrosomes (Figure 3D; Video S7).

Nevertheless, the presence of beads and CLiPs often appeared to partially disrupt the embryo actomyosin network, and we often observed examples where the MT arrays generated by beads and CLiPs managed to interact with nearby spindles. When this occurred, CLiPs and beads formed spindle poles that behaved very similarly to those formed by the endogenous centrosomes (Figure 3E,F; Video S8, S9). These observations establish that some CLiPs functionally resemble centrosomes: they can organise MTs and actomyosin, form spindle poles, and undergo several rounds of duplication—all in synchrony with the embryonic cell cycle.

### Beads coupled to any of the core structural centriole assembly proteins can organise MTs and generate CLiPs

We next asked whether the ability to organise MTs and generate CLiPs was unique to the Sas-6/Ana2 combination, or if it represented a more general property of the structural centriole assembly proteins. To test this, we injected GFP-Nanobody beads into embryos expressing Jup-mCh together with GFP fusions of each individual protein expressed from their endogenous (e) promoter. eAna2-GFP beads were the only ones that organised MTs and generated CLiPs (Figure 4).

**Figure 4.**
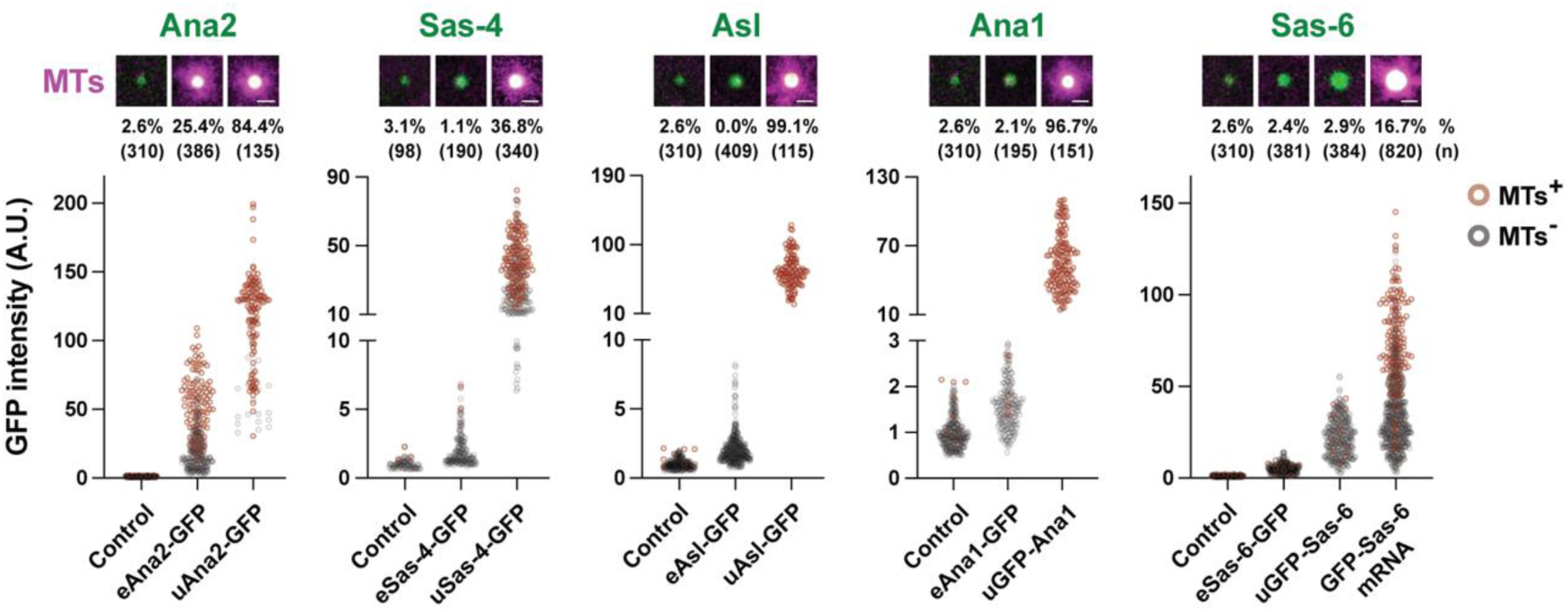
Beads coupled to any one of several structural centriole assembly proteins can organise MTs and generate CLiPs. Images illustrate and graphs quantify the recruitment of MTs (Jup-mCh) (*magenta*) and various GFP-fusions (*green*) to anti-GFP-Nanobody beads injected into embryos expressing the GFP-fusions from either their endogenous (e) or the ubiquitin (u) promoter. Sas-6 beads were also analysed in embryos injected with GFP-Sas-6 mRNA to generate even higher levels of expression. Control beads are coupled to anti-ALFA-tag Nanobodies. Each dot on the graph represents a bead, scored as MT +ve (*brown dots*) or -ve (*grey dots*). The percentage of beads organising MTs (%) and the number of beads analysed (n) is indicated; N=5-10 embryos per condition. Scale bars = 2µm.

Ana2 is expressed at higher levels than the other core assembly proteins in embryos ^44^, potentially explaining why only Ana2 beads were functional. Moreover, western blotting revealed that all the GFP-fusion proteins expressed from their endogenous promoters were present at lower levels in these embryos than their corresponding untagged endogenous proteins (Figure S4). We therefore repeated this experiment using GFP-fusions expressed from the ubiquitin (u) promoter, which expressed these fusions at near, or slightly higher than, endogenous levels (Figure S4). Except for Sas-6-GFP, GFP-Nanobody beads injected into all these embryos organised MTs and generated CLiPs (Figure 4; Videos S10-S13). If, however, we increased Sas-6-GFP levels by mRNA injection, Sas-6-coupled beads also organised MTs and generated CLiPs (Figure 4; Video S14). Thus, any one of the core structural assembly proteins can organise MTs and generate CLiPs if recruited to beads at high enough levels.

### Beads generate CLiPs by recruiting Plk4

How do all these proteins programme beads to organise MTs and generate CLiPs? If they use the canonical assembly pathways, then this functionality should require Polo and Plk4, respectively (Figure 1B). As described above, however, beads that directly recruited Polo or Plk4 did not organise MTs or generate CLiPs, (Figure 2A,B; Videos S1, S2)—even though these beads recruited much higher levels of these kinases than beads coupled to structural assembly proteins that support PCM and CLiP assembly (Figure S5). Thus, if the core structural proteins need to recruit Polo and/or Plk4 to organise MTs and generate CLiPs, they must do so in a specific manner.

We first asked if Plk4 was required to generate CLiPs by examining Sas-6/Ana2 beads in embryos depleted of Plk4 by RNAi. Here, the endogenous centrioles quickly stopped duplicating, but embryos continued to cycle. The Sas-6/Ana2 beads continued to generate MT arrays in synchrony with the remaining endogenous centrosomes, but they no longer generated CLiPs (Figure 5A; Video S15). Thus, Plk4 is required for beads to generate CLiPs, but not to organise MTs.

**Figure 5.**
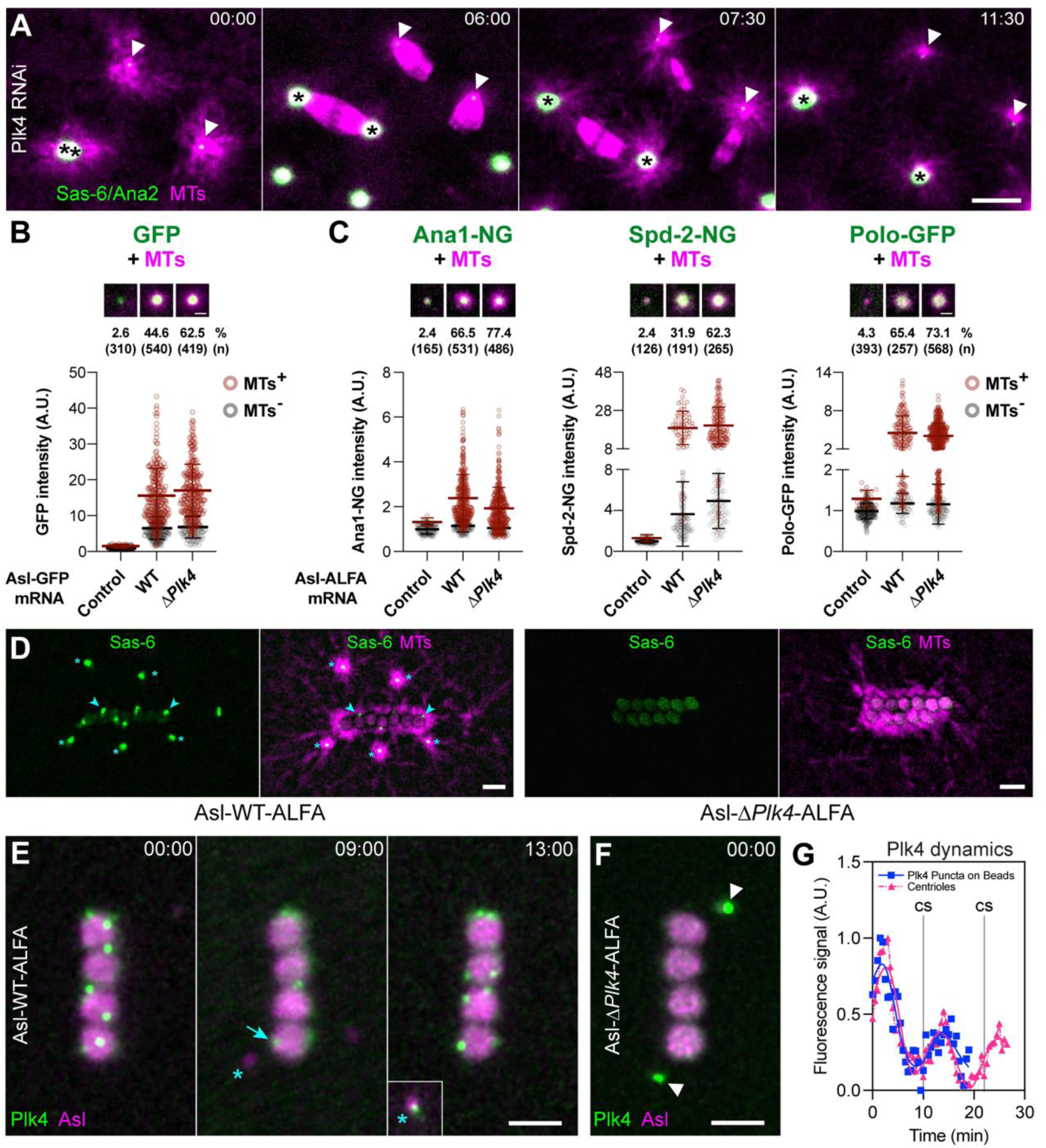
Asl recruits Plk4 to beads to stimulate CLiP assembly. (**A**) Images illustrate the behaviour of Sas-6/Ana2 beads (*black asterisks*) in embryos depleted of Plk4 by RNAi. The beads do not generate CLiPs, although they still organise MTs and can form spindle poles. (N=5 embryos; n≥80 beads analysed). (**B,C**) Images illustrate and graphs quantify the recruitment of Asl-WT-GFP or Asl-Δ*Plk4*-GFP (*green*) to GFP-Nanobody beads (B) or of Ana1-NG, Spd-2-NG or Polo-GFP (*green*) to ALFA-Nanobody beads coupled to either Asl-WT-ALFA or Asl-Δ*Plk4*-ALFA (C) in embryos injected with the indicated mRNA and expressing Jup-mCh (MTs) (*magenta*). Control beads are coupled to the experimental Nanobody and injected without any mRNA. N=7-25 embryos per condition. (**D**) Images illustrate how Asl-WT-ALFA and Asl-Δ*Plk4*-ALFA beads organise MTs (*magenta*), but only the Asl-WT beads generate CLiPs (*cyan asterisks*)—detected here with Sas-6-GFP (centriole marker, *green*). *Cyan arrowheads* highlight some of the Sas-6 foci localised on the surface of beads; CLiPs (*cyan asterisks*) emerge from these foci. N=5-7 embryos; n≥100 beads analysed. (**E**) Images illustrate Plk4-NG dots on the surface of Asl-WT-ALFA beads. The dot highlighted (*cyan arrow*, t=09:00) generates a CLiP (*cyan asterisk*), which later acquires Plk4 (inset, t=13:00; inset shown as this CLiP had moved away from the beads). (**F**) Asl-Δ*Plk4*-ALFA beads do not organise Plk4 dots (the structures highlighted with *white arrowheads* are centrioles in the cytoplasm). (**G**) Graph quantifies Plk4-NG fluorescence levels on bead-associated dots (*blue* squares) and endogenous centrioles (*pink* triangles), illustrating how Plk4 is recruited to both structures in a pulsatile manner. Each line indicates a smoothed spline fit with 8 knots. The grey lines mark the time of centriole separation (CS). Graphs show the mean intensity of 5-6 centrioles/particles imaged every 30secs in a representative embryo. Scale bars = 10μm (A) and 2µm (B-F).

We next asked if Plk4 must be recruited to beads specifically by Asl/Cep152, which normally recruits Plk4 to centrioles. Plk4 interacts with the N-terminal region of Asl/Cep152 ^45–48^, and AlphaFold3 predicted a high confidence interaction between *Drosophila* Asl_18-54_ and Plk4 (ipTM=0.74; Figure S6). We therefore injected ALFA-Nanobody beads together with mRNA encoding either WT Asl-ALFA or a mutant lacking this Plk4-binding region (Asl_ΔPlk4_) into embryos expressing Sas-6-NeonGreen (Sas-6-NG) as a centriole marker and Jup-mCh. WT-Asl-GFP and Asl_ΔPlk4_-GFP were recruited to beads at similar levels (Figure 5B), and WT-Asl-ALFA and Asl_ΔPlk4_-ALFA beads both recruited similar amounts of Ana1, Spd-2 and Polo (Figure 5C). Both sets of beads organised robust MT arrays in synchrony with the cell cycle, but only WT-Asl beads generated CLiPs (Figure 5D; Video S16). Thus, Plk4 must be recruited to beads by Asl to generate CLiPs.

One possible reason for this is that Plk4 presumably must break symmetry on the centriole surface to generate CLiPs. We recently showed that this symmetry breaking could be described as a Turing-like reaction-diffusion system dependent on local interactions between Asl/Cep152 and Plk4 ^49^. This system can theoretically be established on any suitably permissive surface, and the model predicts that larger surfaces should generate more Plk4 foci—potentially explaining why these large 1μm beads frequently produce multiple CLiPs. In support of this model, we observed that 0.5μm beads tended to generate less CLiPs than the 1μm beads (Figure S7).

Plk4-NG did appeare to break symmetry on WT-Asl-ALFA beads as it was highly concentrated in a small number of “dots” on their surface, and these dots functioned as sites of CLiP assembly (*cyan asterisk*, Figure 5E; Video S17). Asl_ΔPlk4_ beads did not organise Plk4-NG dots (Figure 5F; Video S17—the dots highlighted with *arrowheads* are endogenous centrioles). Moreover, in *Drosophila* embryos Plk4 is recruited to centrioles in a pulsatile manner ^50^, and this was also the case on the WT-Asl beads (Figure 5E,G; Video S17).

### Beads organise PCM and MTs by recruiting Polo

These experiments demonstrate that Asl beads generate CLiPs by recruiting Plk4. But Asl beads also organise MTs, even though Asl does not appear to directly recruit Polo/Plk1 to stimulate PCM assembly ^51^ (Figure 1B). Asl beads can recruit Ana1, Spd-2 and Polo (Figure 5C), however, potentially explaining how they engage the PCM assembly pathway to organise MTs (Figure 1B). Moreover, like Asl beads, Ana1 beads organise MTs and generate CLiPs and these beads recruited Spd-2 and Polo (as expected from the canonical PCM assembly pathway) but also Asl, potentially explaining how these beads also generate CLiPs (top panels, Figure 6A). In contrast, Spd-2 beads can organise MTs ^38^, but they do not generate CLiPs (Figure S8), and we found that these beads could recruit Polo (to stimulate the PCM assembly pathway), but not Ana1 or Asl (bottom panels, Figure 6B), presumably explaining why they can organise MTs but cannot generate CLiPs.

**Figure 6.**
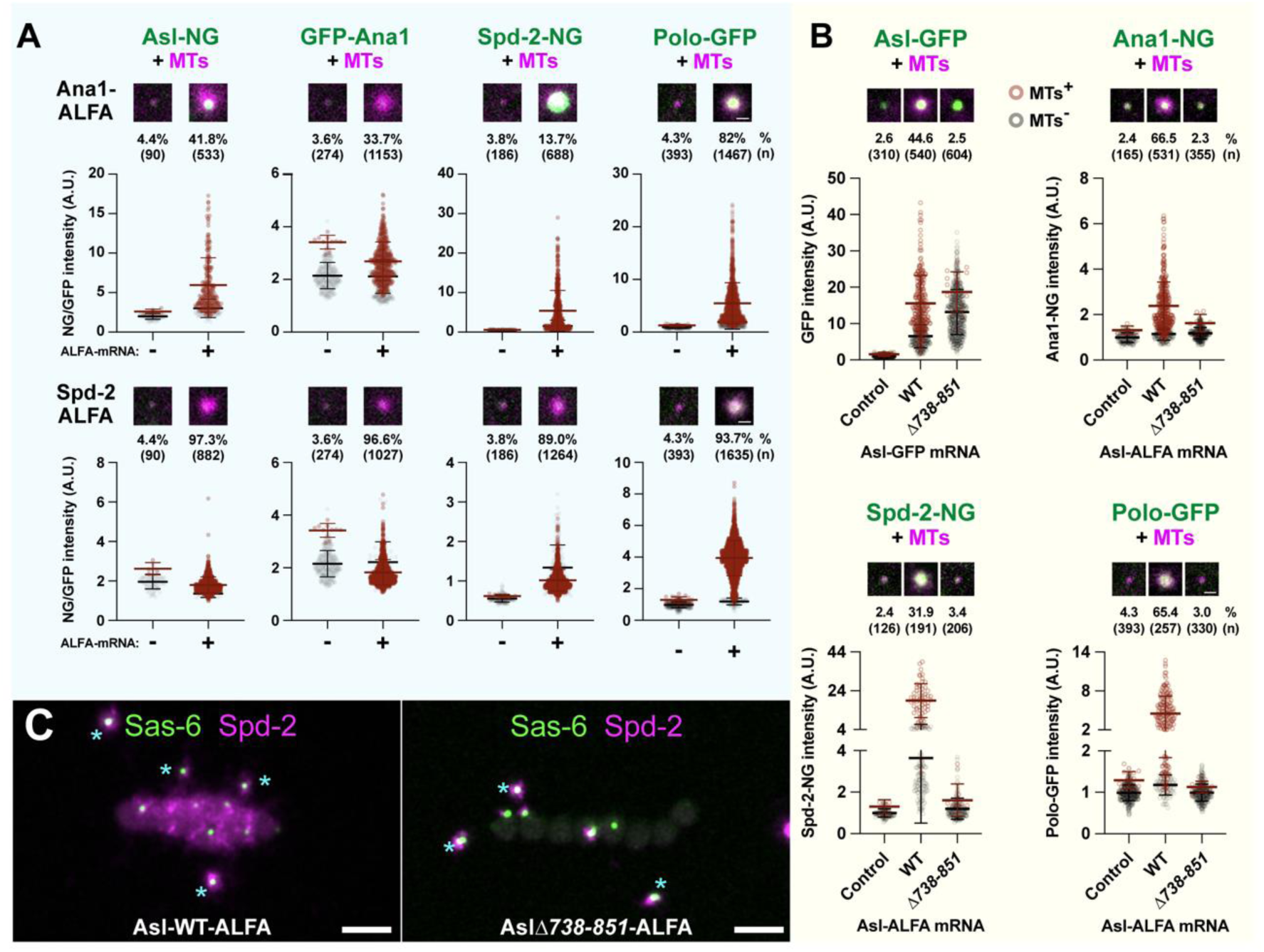
Asl recruits Ana1 and Spd-2 to beads to organise PCM and MTs. (**A**) Images illustrate and graphs quantify the recruitment of Asl-NG, GFP-Ana1, Spd-2-NG or Polo-GFP (*green*) to anti-ALFA-Nanobody beads in embryos expressing Jup-mCh (MTs) (*magenta*) and injected with mRNA encoding ALFA-tagged Asl, Ana1 or Spd-2. Control beads are coupled to the anti-ALFA-Nanobody and injected without any mRNA. The percentage of beads organising MTs (%) and the number of beads analysed (n) is indicated; N=5-10 embryos per condition. (**B**) Images illustrate and graphs quantify the recruitment of Asl-WT-GFP or Asl-*Δ738-851*-GFP to anti-GFP-Nanobody beads (top left), or of Ana1-NG, Spd-2-NG or Polo-GFP (*green*) to anti-ALFA-Nanobody beads in embryos expressing Jup-mCh (*magenta*) injected with mRNA encoding either Asl-WT-ALFA or Asl-*Δ738-851*-ALFA. Control beads are coupled to the appropriate Nanobody and injected without any mRNA. N=7-25 embryos per condition. (**C**) Super-resolution images show Asl-WT-ALFA or Asl*-Δ738-851*-ALFA beads in embryos expressing Sas-6-NG (centriole marker, *green*) and Spd-2-mScarlet-I3 (*magenta*). The Asl-WT beads recruit Spd-2 and generate Sas-6-positive CLiPs (*cyan asterisks*), whereas Asl-Δ*738-851* beads generate CLiPs but do not recruit Spd-2 to their surface. N=3 embryos; n≥20 beads analysed per condition. Scale bars = 2 µm.

We wanted to test the potential importance of the interactions between Asl, Ana1 and Spd-2 in allowing beads to both organise MTs and generate CLiPs. We found that FLAG-Asl co-immunoprecipitated GFP-Spd-2 in S2 cells, and this interaction was disrupted by the internal deletion of Asl_738–851_ (Figure S9), suggesting that this region of Asl either interacts directly with Spd-2, or with Ana1, which then in turn interacts with Spd-2. Asl_Δ738–851_-GFP was recruited to beads as efficiently as WT Asl-GFP (top left panel, Figure 6B), but Asl_Δ738–851_-ALFA beads recruited much less Spd-2, Ana1 and Polo than WT Asl-ALFA beads, and they did not organise MTs (Figure 6B). Strikingly, however, these beads continued to generate CLiPs without organising PCM (Figure 6C; Video S18). Thus, the Asl_738–851_ deletion disrupts the ability of Asl to recruit Ana1 and Spd-2 to engage the Polo-dependent PCM assembly pathway while leaving intact its ability to engage the Plk4-dependent centriole assembly pathway.

Our interpretation of these experiments relies on our assumption that, as we have shown at centrioles, Ana1 recruits Polo to beads to initiate PCM recruitment ^51^ (Figure 1B). To test this, we used a form of Ana1 that cannot recruit Polo (Ana1_ΔPolo_)— previously referred to as Ana1-ALL ^51^. Ana1_ΔPolo_-GFP was recruited to beads as efficiently as WT Ana1-GFP (Figure S10A). But, as expected, Ana1_ΔPolo_-ALFA beads recruited very little Spd-2-NG or Polo-GFP, and failed to organise MTs (Figure S10B), confirming that Ana1’s ability to recruit Polo is essential for this functionality at beads.

### Beads coupled to centriole cartwheel components Sas-4, Ana2 and Sas-6 generate CLiPs and organise MTs by recruiting Asl and Ana1

These experiments indicate how reciprocal interactions between Asl and Ana1 (and potentially Spd-2) allow these beads to both organise MTs and generate CLiPs. It was unclear, however, how beads coupled to the inner cartwheel proteins Sas-6, Ana2 or Sas-4 do this. We suspected that Sas-4 would be crucial, as it links the inner and outer regions of the centriole by binding to both Ana2/STIL ^52,53^ and Asl/Cep152 ^54–56^.

Moreover, *Drosophila* Sas-4 is phosphorylated by CDK/Cyclin at T_200_, generating a Polo-Box binding motif that recruits Polo/PLK1 to daughter centrioles “converting” them to mothers that recruit Asl, and so can recruit PCM and duplicate ^57^. To test whether Sas-4 beads rely on these conversion events to organise MTs and generate CLiPs, we coupled beads to Sas-4 mutants carrying single amino acid substitutions that either prevent the phosphorylation of T_200_ by CDK (T_200_A, P_201_G) or the subsequent recruitment of Polo (S_199_G, S_199_T) ^57^. GFP-fusions of these mutants were recruited to beads as efficiently as WT Sas-4-GFP (Figure S11A), but ALFA-tagged versions did not recruit Polo or Asl, and they did not organise MTs or generate CLiPs (Figure S11B). Thus, remarkably, a single, highly conservative, S_199_T substitution blocks Sas-4’s ability to stimulate MT organisation and centriole/CLiP generation at both centrioles and beads. Moreover, we found that the ability of Ana2 beads to organise MTs and generate CLiPs depended on its recruitment of Sas-4, but not on its interaction with itself or with Sas-6 (Figure S11D-H). Finally, Sas-4, Sas-6 and Ana2 beads could all recruit Asl, Ana1 and Spd-2, but Sas-4 beads did not need to recruit Sas-6 or Ana2 to do so (Figure S12). We conclude that beads bound to the inner cartwheel proteins organise MTs and generate CLiPs because they ultimately recruit Sas-4 to engage the canonical assembly pathways (Figure S13).

### CLiPs share several architectural hallmarks of centrosomes

Our data so far indicates that CLiPs are functionally similar to centrosomes and are built using the canonical assembly pathways. We therefore wanted to test whether CLiPs shared a common architecture with centrosomes. We first compared the distribution of NG-Asl at endogenous centrosomes and CLiPs as they separated at the end of mitosis (*orange* and *cyan* boxes, respectively, Figure 7A). At this stage, old and new mother centrioles often maintained their orthogonal orientation, providing simultaneous end-on (*arrowheads*, Figure 7A) and side-on (*arrows*, Figure 7A) views of the Asl torus that surrounds the mother centriole when viewed by super-resolution microscopy. Separating CLiPs assembled a similar Asl torus, recapitulating this distinctive structural feature of centrioles (Figure 7A; Video S19). Quantification, however, revealed that the Asl torus was slightly smaller at CLiPs than at centrioles (Figure S14A), and that the fluorescence levels of several centrosome markers were slightly dimmer (Figure S14B). This is likely because, as described above, some CLiPs are not fully mature at birth.

**Figure 7.**
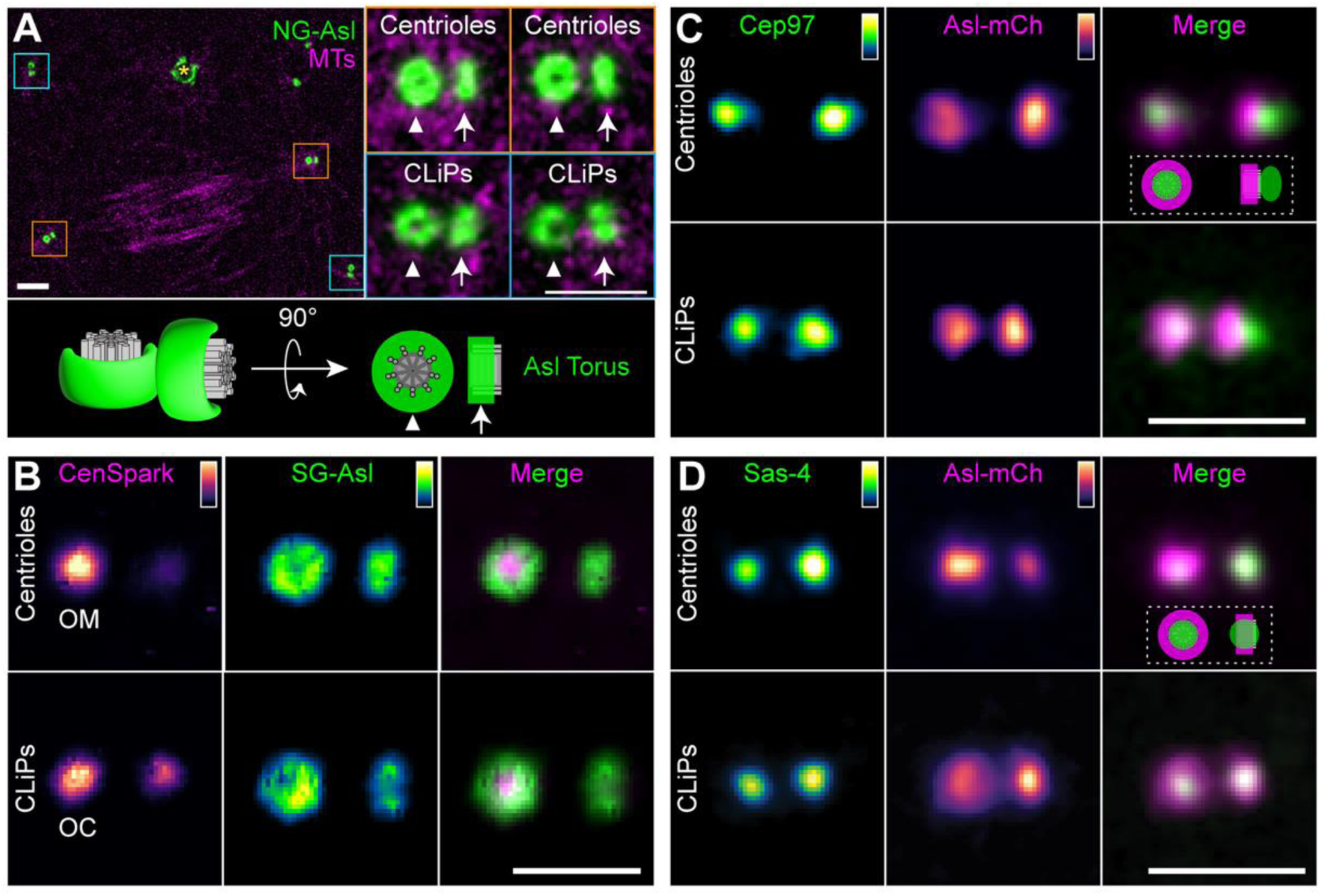
CLiPs share several structural features with centrosomes. (**A**) Two-colour super-resolution images of NG-Asl (*green*) and MTs (Jup-mCh) (*magenta*) in a living embryo highlighting the duplicated centrioles separating at the end of mitosis at the poles of a spindle (*orange boxes*) and two duplicated CLiPs separating at the same time in the cytoplasm (*cyan boxes*). A bead in the embryo is indicated with a *yellow asterisk*. Magnified insets, and illustration below, highlight the top-down barrel (*arrowhead*) or side-on bar (*arrow*) view of the Asl torus on the centrioles/CLiPs. The magnified insets have been rotated to align their orientations. (**B-D**) Two-colour super-resolution images of duplicated centrioles/CLiPs separating at the end of mitosis in embryos expressing various centriole markers (as indicated, see main text for details). Asl is not clearly visualised as a torus in (C,D) because the C-terminal Asl-mCh tag is located closer to the centrioles than the N-terminal NG- or SG-Asl tags shown in (A,B). All panels show representative images: N=4-6 embryos; n=20-40 centrioles/CLiPs analysed in total. Scale bars = 1µm.

We next asked whether CLiPs contain MT doublets or triplets—a unique and defining feature of centrioles. The fluorescent small molecule CenSpark specifically labels MT doublets/triplets ^58^, so we co-injected it into embryos that were generating CLiPs from Asl-ALFA beads and expressing StayGold-Asl (SG-Asl). As expected, CenSpark labelled the endogenous centrioles internally to the Asl torus, and a similarly positioned structure in CLiPs (Figure 7B). Strikingly, CenSpark more strongly labelled the older centriole/CLiP, suggesting that in these rapidly dividing embryos MT doublets are not fully assembled on new centrioles/CLiPs as they separate from their mothers. This relatively late acquisition of CenSpark labelling could also be observed on CLiPs as they assembled at, and then separated from, the bead surface (Video S20).

Centrioles also have a proximal-distal polarity, and the distal-end capping protein Cep97-GFP was positioned distally to the more proximal Asl-mCherry torus in both centrioles and CLiPs (Figure 7C). In contrast, the cartwheel protein Sas-4-StayGold (Sas-4-SG) colocalised with Asl-mCherry towards the more proximal-end of centrioles and CLiPs (Figure 7D). Finally, nine other centriole/centrosome proteins that we tested exhibited very similar distinctive distributions at centrosomes and CLiPs (Figure S15). Together these data suggest that centrosomes and CLiPs share several architectural features.

## Discussion

Since the earliest observations that centrosomes, like chromosomes, are faithfully transmitted from one cell generation to the next, the question of how they replicate has been one of the great mysteries in cell biology ^59^. Unlike chromosomes, whose replication mechanism was immediately suggested by the discovery of the DNA double helix, our increasing knowledge of centrosome architecture has provided little insight. If anything, the extraordinary molecular and structural complexity of centrioles has only deepened the mystery: how can such elaborate structures duplicate with such precision, and where does the information that guides their assembly reside?

It is well established that centrioles can assemble *de novo* in certain developmental and experimental contexts, demonstrating that a pre-existing mother centriole template is not required for centriole biogenesis ^11–14,16^. Nevertheless, canonical centriole duplication in proliferating cells remains highly dependent on a pre-existing mother centriole. When *de novo* assembly is induced experimentally in proliferating cells, the process is typically slow and error prone ^17–22^. In this context, our observation that a synthetic bead—programmed to recruit just one protein—can direct the assembly of complex centrosome like particles (CLiPs) in synchrony with the rapidly duplicating endogenous centrosomes is remarkable.

These synthetic beads organise MTs and generate CLiPs because they engage the canonical centriole and PCM assembly pathways (Figure S13). Unlike the structurally and molecularly elaborate mother centriole, however, beads provide a minimal and programmable surface, allowing the function of individual assembly proteins to be examined in isolation. We find that beads coupled to any one of the five core structural proteins that are essential for centriole duplication in flies—Sas-6, Ana2/STIL, Sas-4/CPAP, Asl/Cep152, or Ana1/Cep295—can programme beads to recruit PCM and to generate CLiPs if they are recruited to beads at high enough levels. These multiple inputs seem to converge on the recruitment of Asl/Cep152: Asl recruits Plk4/PLK4 to initiate centriole assembly, and Ana1 and/or Spd-2 to recruit Polo/PLK1 to initiate PCM assembly. In an accompanying paper, Aich and Gönczy show that a similar logic appears to operate in human cells, where Cep63 can recruit Cep152/Asl and Plk4 to engineered condensates to seed centriole cartwheel assembly ^60^.

The ability of Asl and Ana1 to recruit each other to beads probably explains why beads coupled to any one of the core centriole assembly proteins can generate both MTs and CLiPs. This reciprocal recruitment may also explain why the ability of centrioles to assemble PCM and to duplicate appears to be so intimately linked ^61–64^. This intimate association has widely been interpreted as indicating that the PCM helps support centriole assembly. An important conclusion from our work, however, is that these processes can be separated. Asl beads that cannot recruit Plk4 cannot generate CLiPs but can still recruit PCM; Asl beads that cannot recruit Ana1 and/or Spd-2 cannot recruit PCM but can still generate CLiPs.

Perhaps more importantly, we find that the direct recruitment of Plk4 or Polo to the bead surface is insufficient to initiate CLiP or PCM assembly, despite producing much higher local kinase concentrations than recruitment to beads via the other assembly proteins. This suggests that the local environment created by interactions between the kinases and their recruiting proteins is required to stimulate assembly. A crucial insight from our experiments is that establishing this permissive environment is an intrinsic property of these proteins, rather than of any specific feature of the mother centriole.

### Seed—Scaffold—Self-Assemble: a general paradigm for organelle biogenesis?

Our observations provide a surprisingly simple answer to the question of how cells build these elaborate structures, and where the information to do so comes from. Localised kinase activation, coupled with the local concentration of a small number of assembly factors, acts as a seed that initiates the assembly of a relatively simple scaffold (comprising Sas-6, Ana2/STIL and Sas-4/CPCP for centriole assembly, and Spd-2, Cnn and TACC for PCM assembly). These scaffolds then guide the robust self-assembly of centrioles and PCM. In this scenario, the information to guide centrosome assembly resides largely within the centrosome building blocks themselves.

Protein self-assembly is a fundamental organising principle in biology, but it is perhaps surprising that it might explain the assembly of centrosomes—as these organelles are composed of tens to hundreds of copies of more than a hundred different proteins ^10,61–64^. Importantly, the embryo provides at least one additional source of information to guide assembly—the embryonic cell cycle oscillator. Centrosome duplication is tightly coordinated with the cell cycle, and in the early *Drosophila* embryo the cell cycle oscillator is spatially organised around the endogenous nuclei and centrosomes ^65,66^. This organisation presumably explains why synthetic beads do not generate CLiPs until endogenous centrosomes and nuclei arrive at the embryo cortex, thus exposing the beads to oscillations in cell cycle activity for the first time. By creating alternating permissive (S-phase) and non-permissive (M-phase) environments, the embryonic oscillator entrains centrosome and CLiP biogenesis with other cell cycle events. Whether the oscillator provides additional instructive information during the assembly process is unknown.

Our findings therefore define a simple **Seed–Scaffold-Self-Assemble** design logic for centrosome biogenesis. In this framework, a molecular *seed* locally concentrates enzymes and a small number of assembly factors to form a *scaffold* that then initiates the *self-assembly* of a much larger molecular structure. This presumably explains why centrioles can form *de novo*: a molecular seed—likely to contain Asl/Cep152 and Plk4—presumably initiates this process ^15,16,64^. It can also explain centriole biogenesis in multiciliated cells, where specialised organelles (deuterosomes) that lack the highly ordered structure of centrioles—but that contain Asl/Cep152 and PLK4—act as seeds to generate multiple centrioles from their surface ^67^.

We speculate that similar principles may underlie the biogenesis of many other cellular assemblies, including biomolecular condensates, kinetochores, and membrane-bound organelles that arise at specialized contact sites ^68–71^. Seed–Scaffold-Self-Assemble therefore provides a parsimonious framework for understanding organelle biogenesis more broadly, explaining how relatively simple molecular inputs can generate highly complex cellular structures without the need to directly copy a pre-existing template. An obvious advantage of this framework is that the assembly and activation of the seeds that drive organelle assembly can be tightly spatially and temporally regulated by multiple cellular inputs (such as the cell cycle oscillator in the case of centrosomes). Defining how molecular seeds establish and modulate the reaction-diffusion landscapes that guide self-assembly will be an important challenge for future work. Understanding these principles may ultimately enable the rational engineering of complex synthetic self-assembling nanomachines built from multiple copies of hundreds of different proteins.

### Limitations of the study

Although we show that CLiPs and centrosomes can share many functional and architectural features, it is not clear that they are identical. CLiPs are centred around a structure that is labelled by CenSpark (which specifically labels centriole MTs), but we do not know whether these MTs recapitulate the characteristic 9-fold symmetrical arrangement of *bona fide* centrioles. EM or Expansion Microscopy approaches are technically very challenging in the early embryo system and, even if nine-fold symmetric MT structures were observed, this would not prove that CLiPs and centrosomes are identical. In an accompanying manuscript, Aich and Gönczy use a related strategy to reconstitute centriole biogenesis on an engineered platform in human cells and find that these structures are 9-fold symmetric ^60^. Further work will be required to determine if this is also the case in *Drosophila* CLiPs.

Finally, centrioles organise two important cellular organelles, centrosomes and cilia. Early fly embryos do not form cilia so we cannot assess whether CLiPs have this functionality. Because the CLiPs are not associated with nuclei, they are internalised into the yolk when the embryo cellularises and so are effectively eliminated from the developing embryo. Thus, although CLiPs appear to be fully functional as centrosomes in the early embryo, we cannot assess whether they would be fully functional in later development.

## Acknowledgements

We thank members of the Raff Laboratory for many helpful discussions and comments.

## Funding

Wellcome Trust Investigator Award 215523 (JWR; supporting CCC, MP, SSW, NM, JMM)

Wellcome Trust Discovery Award 304037 (JWR; supporting CCC, MP, SSW, NM, JMM)

National Science and Technology Council, Taiwan, Overseas Project for Postgraduate Research grant 111-2917-I-564-012 (CCC)

Ministry of Science and Technology, Taiwan, Graduate Students Study Abroad Program 112-2917-I-002-026 (MP)

National Science and Technology Council, Taiwan, Overseas Project for Postgraduate Research grant 114-2917-I-564-008 (MP)

Cancer Research UK Oxford Centre Prize DPhil Studentship C5255/A23225 (SSW) Balliol College Jason Hu Scholarship (SSW)

Clarendon Scholarship (SSW)

Max Planck–Croucher Postdoctoral Fellowship (SSW)

Wadham College Junior Research Fellowship in Medical Sciences (SSW)

European Commission Marie Skłodowska-Curie Individual Fellowship 892857 – CENTROMD (JMM)

European Molecular Biology Organization Long-Term Fellowship LTF 29-2019 (JMM)

## Author contributions

Conceptualization: MP, SSW, JMM, TH, CCC, JWR

Methodology: MP, SSW, BX, LBM, LH, ZF, NM, MH, SS, ZN, CCC, JWR

Investigation: MP, SSW, BX, LBM, CCC, JWR

Resources: MP, SSW, BX, LBM, NM, MH, SS, ZN, CCC, JWR

Data curation: MP, SSW, BX, LBM, CCC, JWR Formal analysis: MP, SSW, CCC, JWR Software: MP, SSW, CCC, JWR

Validation: MP, SSW, BX, LBM, CCC, JWR

Visualization: MP, SSW, BX, LBM, CCC, JWR Project administration: MP, SSW, CCC, JWR Supervision: MP, SSW, CCC, JWR

Writing – original draft: MP, SSW, CCC, JWR

Writing – review & editing: MP, SSW, BX, LBM, JMM, CCC, JWR

## Supplementary Figures

**Figure S1.**
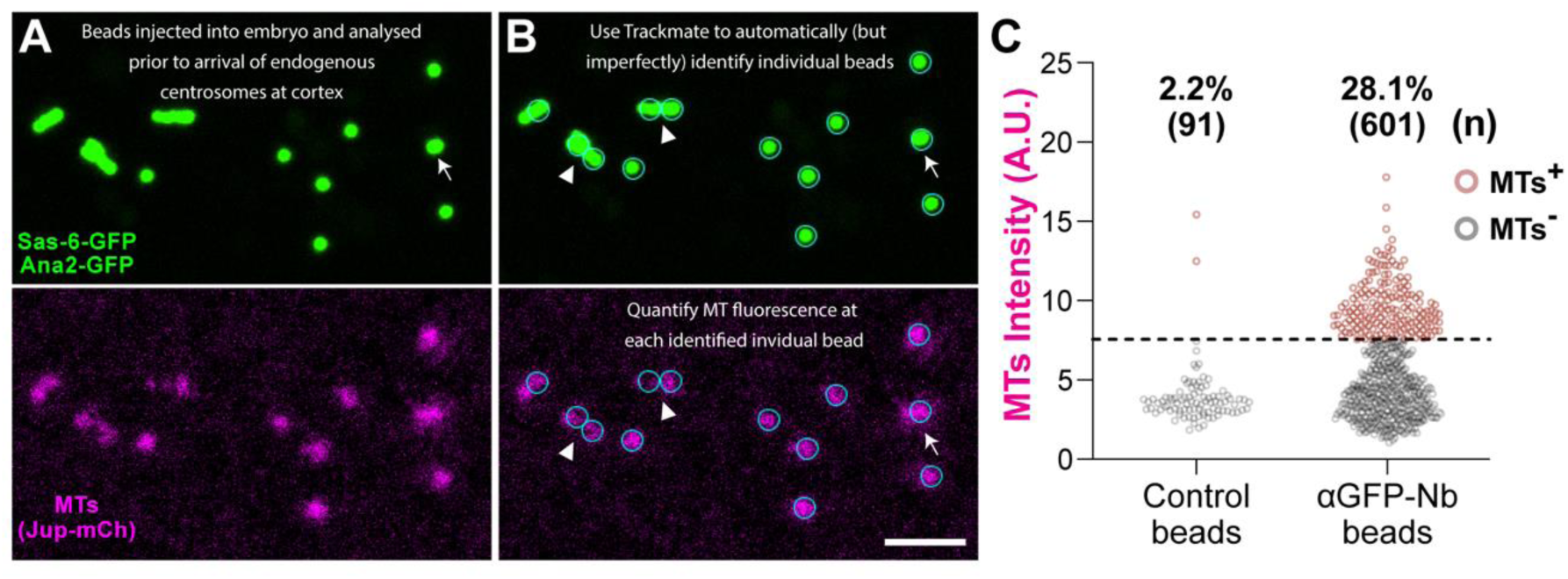
Methodology for automated bead identification and quantification. (**A**) Images show a field of anti-GFP-Nanobody (Nb) beads ∼20-30 mins after injection into an early embryo expressing Sas-6-GFP, Ana2-GFP (*green*) and Jup-mCh (MTs) (*magenta*), before the endogenous centrosomes have arrived at the embryo cortex. In most such embryos there was a region where the beads were well spaced and organised MTs efficiently (as the MTs tend to push the beads apart, *right side*), and another region where the beads tended to clump together and organised MTs less efficiently *(left side*). In these latter regions it was difficult to assess the number of beads and how many organised MTs. (**B**) To quantify this data in an unbiased manner we used the Fiji plug-in Trackmate to detect individual beads based on their GFP-fluorescence (*cyan circles*) or, for Controls, on their weak red-autofluorescence (see Figure 2C). The MT fluorescence intensity of each bead was quantified and plotted on a graph (**C**). Each bead was scored as MT +ve or -ve using a threshold (dashed line) calculated from the Control-bead distribution (Control mean + 2 SD): beads above/below this threshold were scored as +ve (*brown* dots)/-ve (*grey* dots). The percentage of +ve (%) beads and the number analysed (n) are shown. A background +ve rate of ∼1-5% was usually observed in Controls. On inspection, +ve Control beads rarely organised the extensive MT arrays observed at experimental beads, and the occasional bright Control was usually located close to the meiotic spindle remnant. We note that this method is imperfect as bead clusters are often not counted correctly (e.g., the beads highlighted with arrowheads clearly contain more than the two beads identified by TrackMate). As this error occurred consistently in all our analyses, we believe that it did not alter any of our conclusions. Scale bar = 10µm.

**Figure S2.**
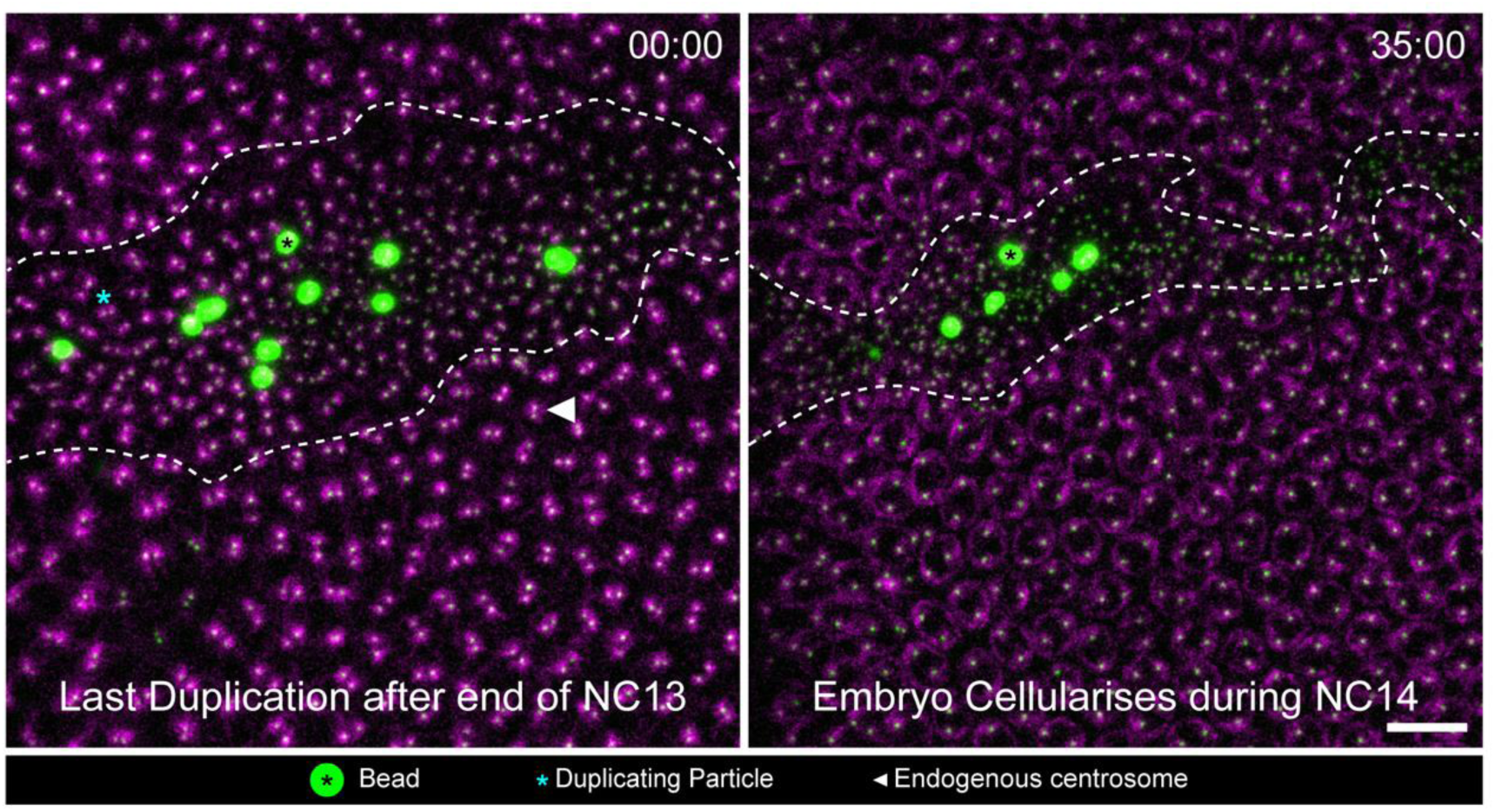
Beads and CLiPs stop duplicating at NC14 and are gradually internalised during cellularisation. Images show Sas-6/Ana2-GFP beads (*black* asterisk) in a living embryo expressing Sas-6-GFP, Ana2-GFP (*green*) and Jup-mCh (MTs) (*magenta*). At the start of NC14 (t=00:00) the endogenous centrosomes that duplicated in NC13 have just separated (*arrowhead*), as have most of the duplicated CLiPs (dotted line delineates the approximate boundary between the endogenous nuclei/centrosomes and beads/CLiPs). At this time the nuclei and centrosomes stop dividing, and the embryo starts to cellularise. As cellularisation proceeds (t=35:00), the beads and CLiPs, which are not associated with any nuclei, are compacted into a gradually shrinking area of the cortex, and they are eventually internalised into the yolk, so they do not contribute further to post-cellular embryonic development. This is a natural process that normally eliminates any cortical centrosomes that have become detached from damaged nuclei ^1^. It was often hard to follow individual CLiPS as they became internalised, but a careful inspection failed to detect any duplication events during this process. N=5 embryos; n≥200 CLiPs analysed. Video S3 shows an embryo undergoing this process, although it is different to the one shown in this Figure. Scale bar = 10 µm.

**Figure S3.**
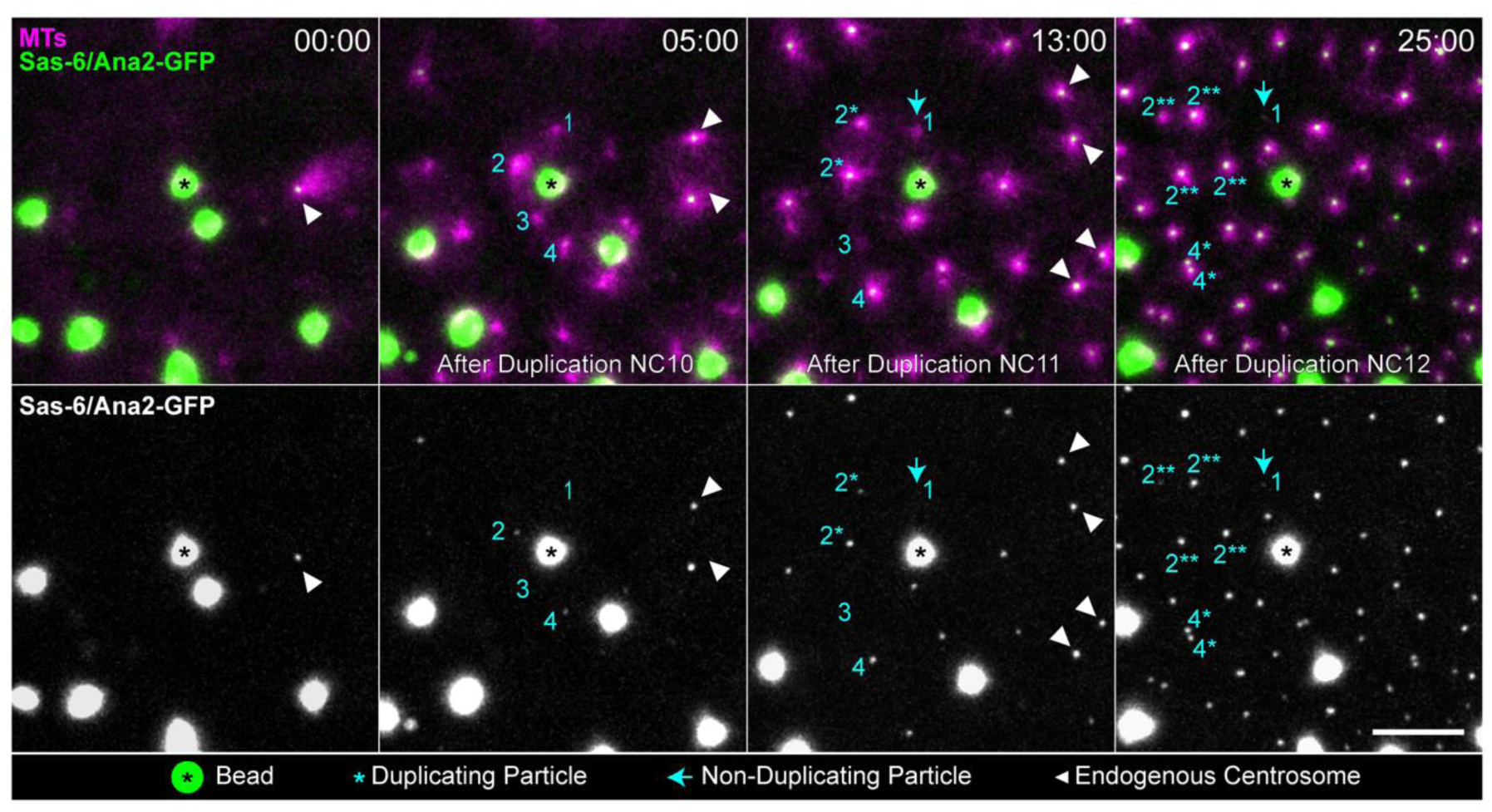
Particles generated by Sas-6/Ana2 beads are born with variable levels of Sas6/Ana2 that seem to dictate their fate. Images show stills from a video of Sas-6/Ana2-GFP beads (*green*) in an embryo expressing Jup-mCh (MTs) (*magenta*), tracked over three successive nuclear cycles (NC10-NC12). The top row shows the merged channels; the bottom row shows the Sas-6/Ana2-GFP signal alone, displayed with increased brightness to visualise dimmer particles that would otherwise be difficult to detect. Although many particles are born with levels of Sas-6/Ana2-GFP that are similar to the endogenous centrrioles and can proceed through multiple rounds of high-fidelity duplication (e.g., Figure 3B; Video S4), the beads shown here generate four particles (numbered 1-4; t=05:00) that we use to illustrate how some CLiPs are born with lower levels of Sas-6/Ana2 and can have different fates. **Particle 2** is born with the highest levels of Sas-6/Ana2 (although lower than the endogenous centrosomes, highlighted with *arrowheads*, t=5:00). This particle initially duplicates to generate two CLiPs, (2*, t=13:00), one of which is dimmer than the other. Both particles duplicate again during the next cycle to generate four CLiPs, (2**; t=25:00), one of which remains dimmer than the others. **Particle 4** is born with slightly lower levels of Sas-6/Ana2 and it does not duplicate during the first round of division (t=13:00). But it accumulates more Sas-6/Ana2 and duplicates in the next cycle (4*, t=25:00). **Particles 1 and 3** organise MTs but are born with essentially undetectable levels of Sas-6/Ana2. These particles are still visible at t=13:00, but by t=25:00 Particle 3 is no longer detectable, while Particle 1 is visible as a tiny dot of Sas-6/Ana2 labelling (*arrow*, t=25:00). Scale bar = 10μm.

**Figure S4.**
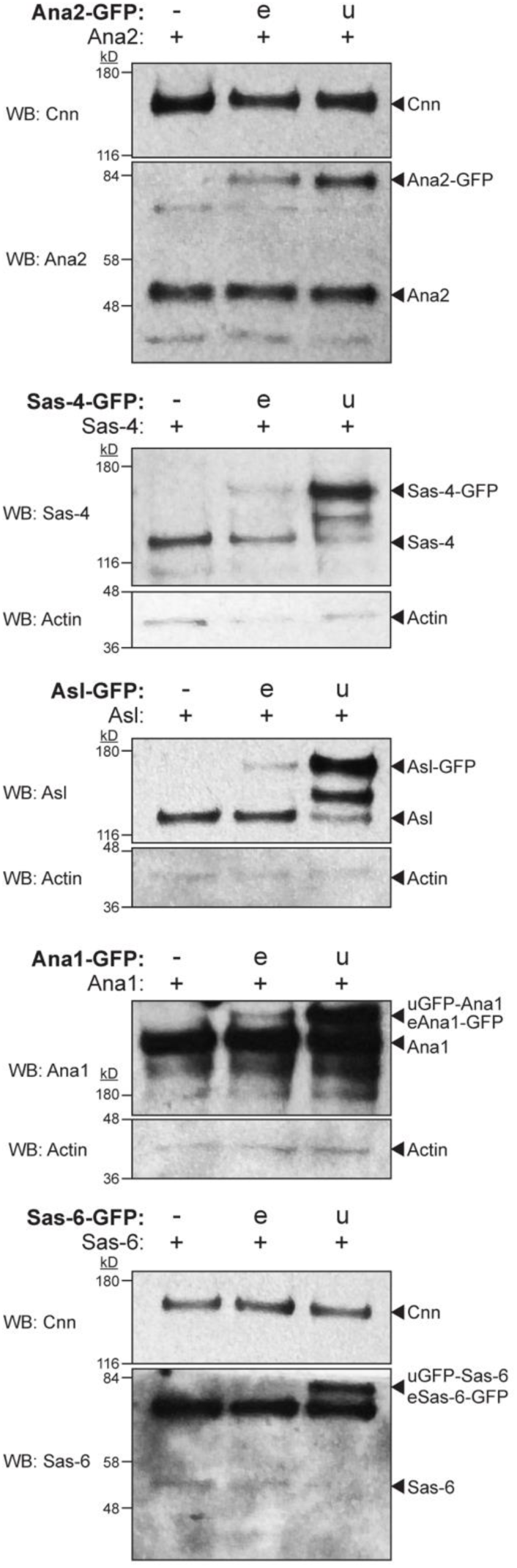
Western blot analysis of endogenous and ubiquitin promoter-driven GFP-fusion expression levels. Western blots of early embryos from transgenic *Drosophila* lines expressing GFP-fusions of Ana2, Sas-4, Asl, Ana1 or Sas-6 from either their endogenous (e) or the ubiquitin (u) promoter. The first lane in each blot (–) shows a WT control expressing no GFP-fusion. Most of these transgenic lines expressed one copy of the transgene and two copies of the endogenous WT gene, but the eAna2, eSas-4, uSas-4 and uAsl lines were recombined with individual mutants and therefore carried only a single copy of the endogenous gene. These were the combinations used in all the experiments reported here, unless otherwise stated. Cnn or Actin were used as loading controls. In all these embryos, the endogenous promoter lines express lower levels of the GFP-fusion compared to the endogenous untagged protein, whereas the ubiquitin promoter drives expression of each GFP-fusion at levels broadly comparable to, or at slightly higher levels than, the endogenous protein. Molecular weight markers (kD) are indicated. Blots are representative examples from N=2 independent experiments.

**Figure S5.**
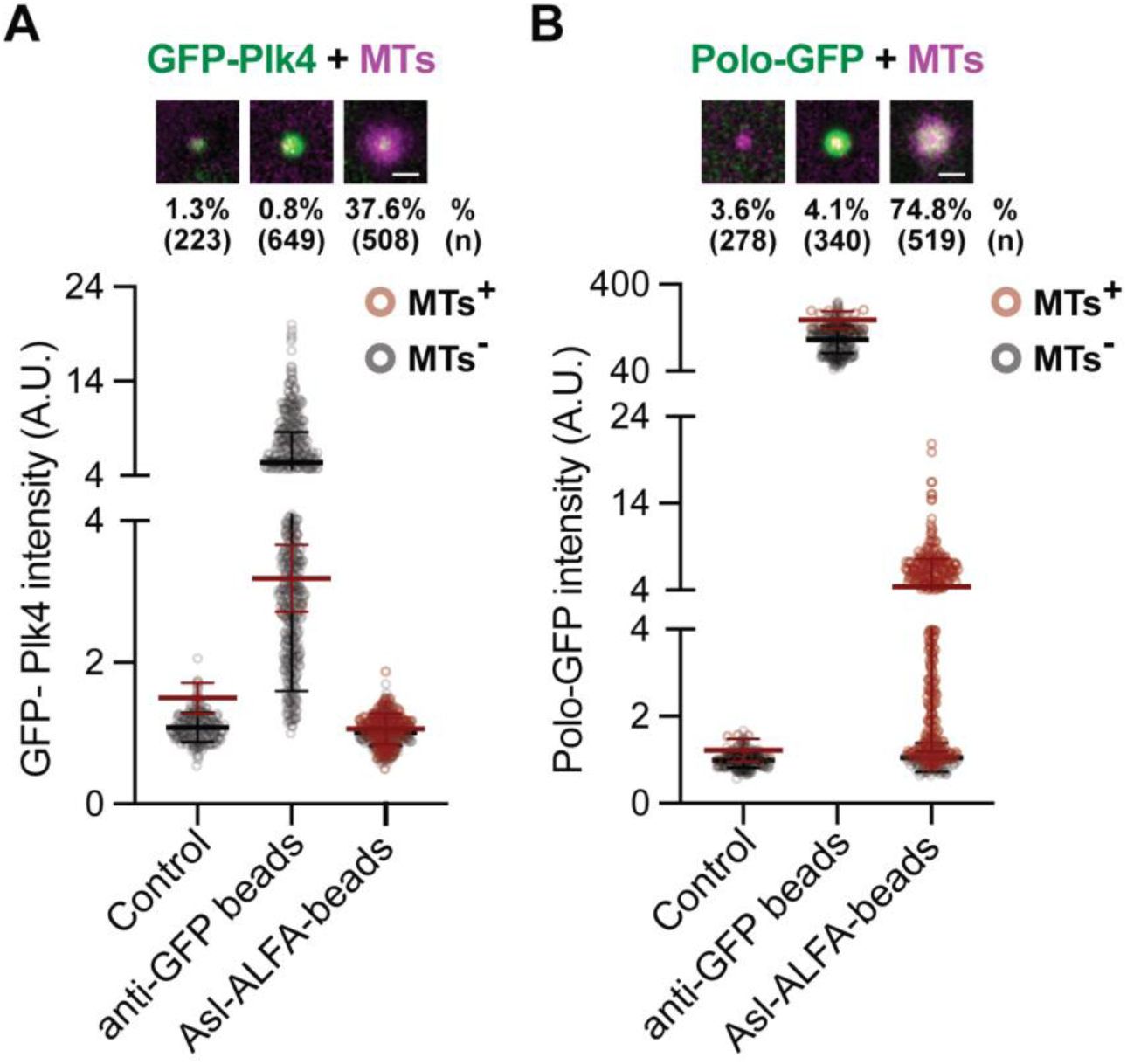
Beads that directly recruit high levels of Plk4 or Polo don’t organise MTs or generate CLiPs, but Asl-ALFA beads with lower levels of either do both. (**A,B**) Images illustrate and graphs quantify the recruitment of GFP-Plk4 (A) or Polo-GFP (B) (*green*) and MTs (Jup-mCh) (*magenta*) at either anti-GFP beads that directly recruit each kinase, or at Asl-ALFA beads that indirectly recruit each kinase via Asl. The percentage of beads organising MTs (%) and the number of beads analysed (n) is indicated; N=4-18 embryos per condition. The anti-GFP-Nanobody beads directly recruit relatively high levels of GFP-Plk4 or Polo-GFP but do not organise MTs or generate CLiPs (see Figure 2A,B). In contrast, Asl-ALFA beads recruit substantially lower levels of Plk4 or Polo yet efficiently organise MTs and generate CLiPs (see Video S12). This suggests that simply concentrating Plk4 or Polo on a bead is not sufficient to nucleate MTs or generate CLiPs; rather, these functions require Asl-dependent recruitment. Scale bars = 2 µm.

**Figure S6.**
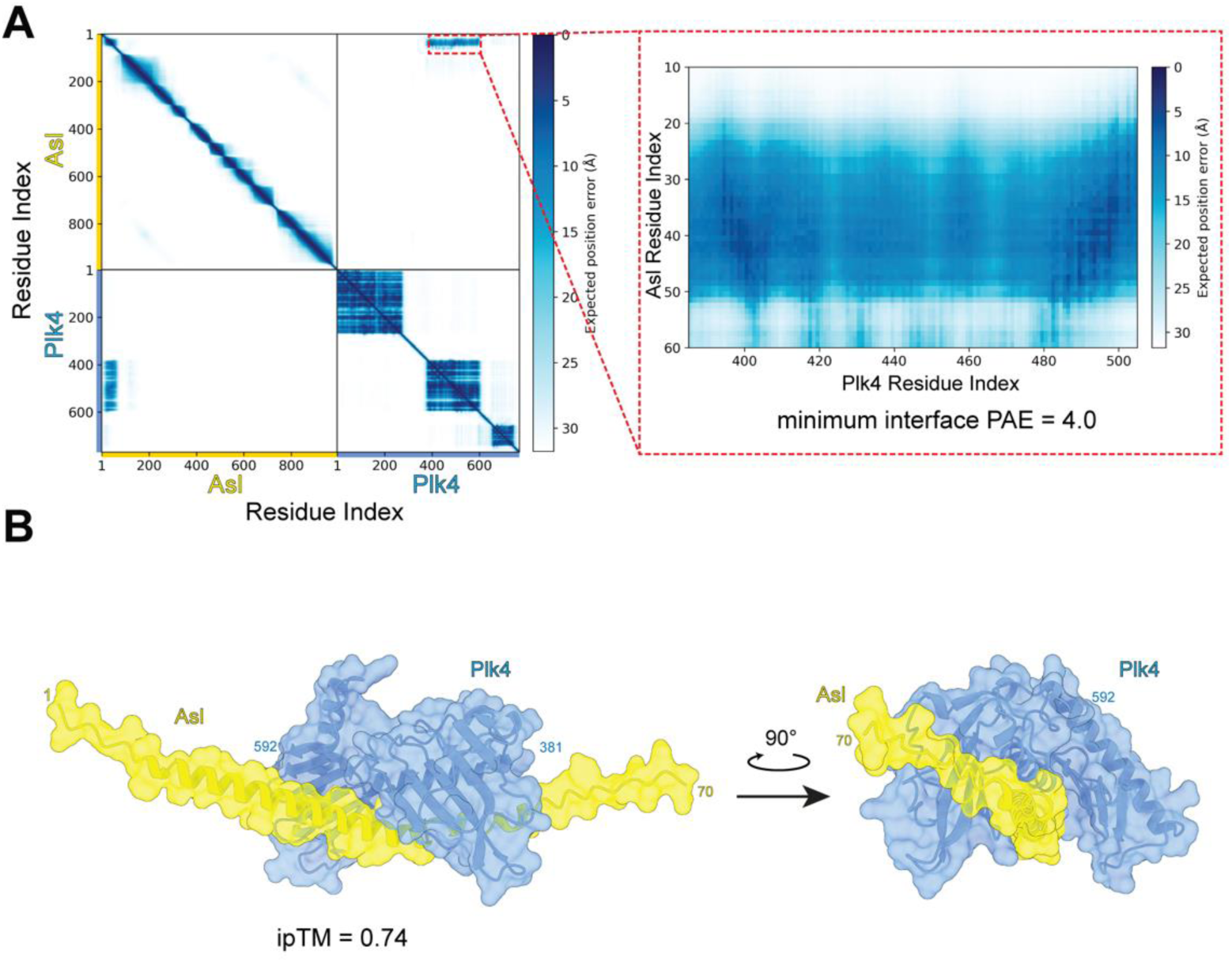
AlphaFold3 predicts a high-confidence interaction between the N-terminus of Asl and the C-terminal region of Plk4. (**A**) Predicted Aligned Error (PAE) plot from an AlphaFold3 prediction of the full-length Asl–Plk4 complex (left). Dark blue indicates low expected position error (high confidence), and off-diagonal low-PAE regions indicate residue pairs whose relative positions are predicted with high confidence—consistent with a direct interaction. The *dashed red box* shows a magnified view of the sole inter-chain interaction interface, between Asl residues ∼10-55 and Plk4 residues ∼380-500, with a minimum interface PAE of 4.0 Å. (**B**) Predicted structural model of the Asl–Plk4 interface, shown in two orientations related by a 90° rotation. Asl (*yellow*) forms an extended helical segment (residues 1-70 shown) that binds along a groove on the surface of the Plk4 C-terminal region (*blue*; residues 381-592 shown). Residue numbers at the visible chain termini are indicated.

**Figure S7.**
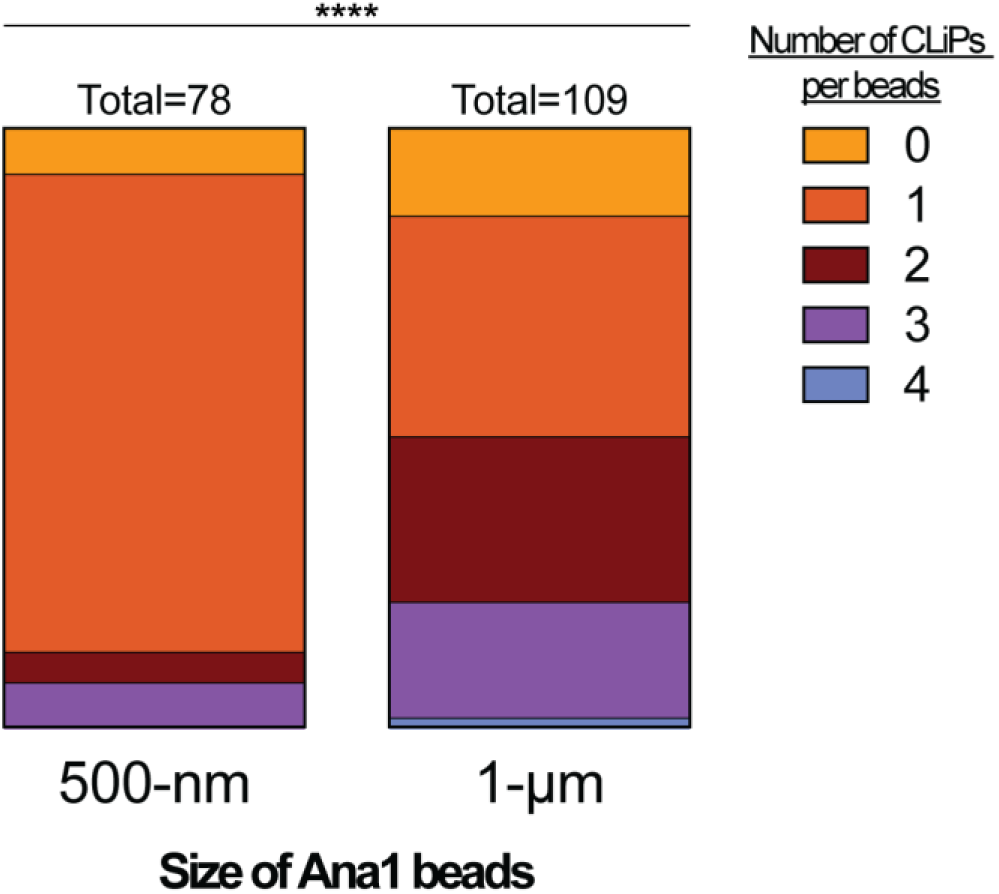
0.5μm beads tend to generate fewer CLiPs than 1μm beads. Bar charts quantify the number of CLiPs generated in a single cycle from GFP-Nanobody beads of different sizes injected into embryos expressing Ana1-GFP and Jup-mCh. In this experiment beads were assessed for whether they generated at least one CLiP in NC11; beads that did so were then assessed for the number of CLiPs they generated at the end of NC12. This experiment shows that the smaller, 0.5μm, beads tend to generate fewer CLiPs than the larger 1μm beads, as predicted by the Turing-type reaction-diffusion model proposed to explain Plk4 symmetry breaking ^2^. A caveat to this experiment is that the 0.5μm beads are made using a different chemistry to the1μm beads (see Star Methods).

**Figure S8.**
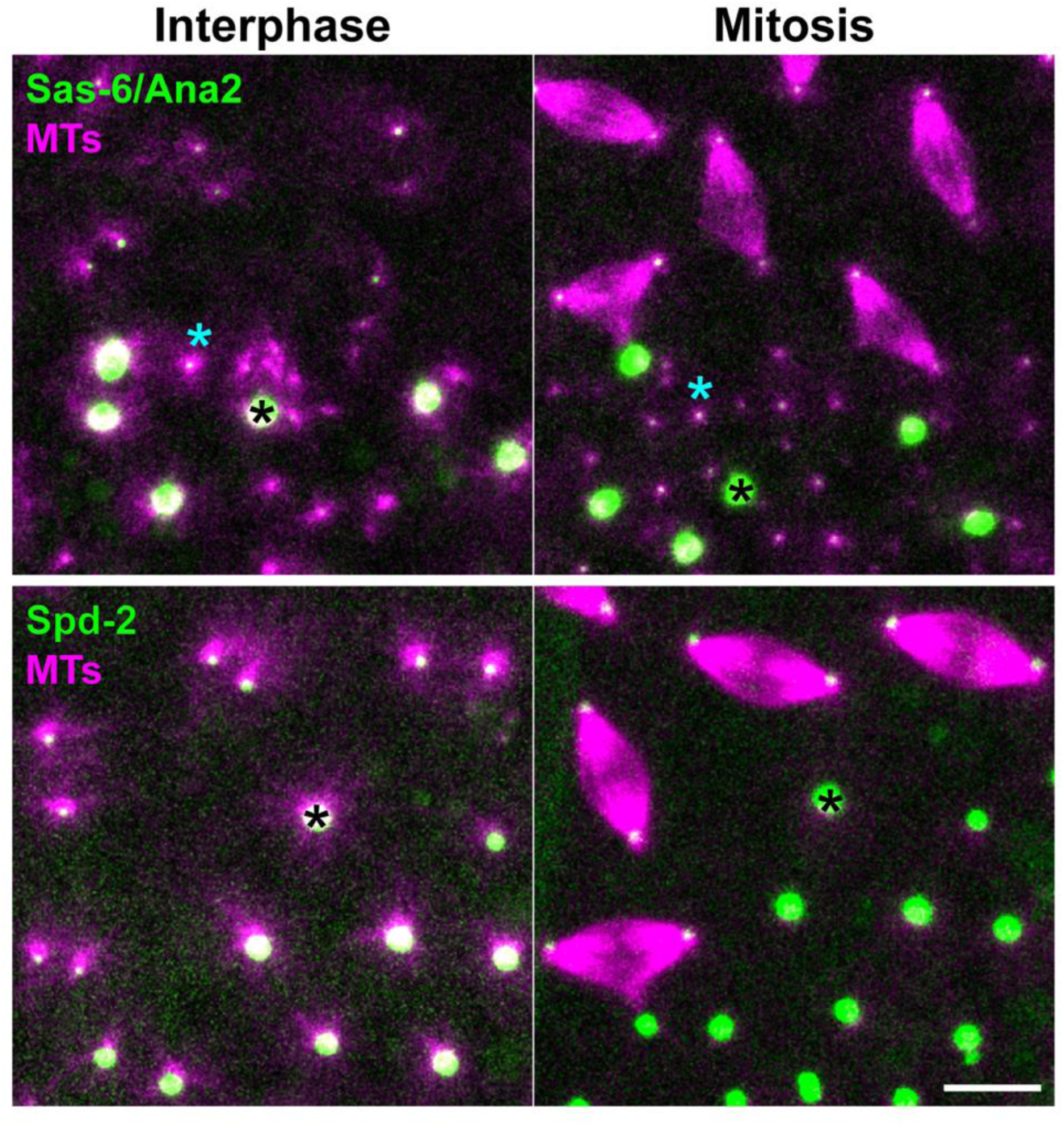
Spd-2 beads organise MT arrays that cycle in phase with the endogenous centrosomes, but they do not generate CLiPs. Images show anti-GFP-Nanobody beads ∼30 mins after injection into embryos expressing either Sas-6-GFP and Ana2-GFP (top) or Spd-2-GFP (bottom) (*green*), together with Jup-mCh (MTs) (*magenta*), imaged during interphase (left) and mitosis (right). Sas-6/Ana2 beads generate CLiPs that separate from the bead surface; like the endogenous centrosomes, the beads and CLiPs organise long MT arrays in interphase and short arrays in mitosis (*black asterisks* highlight a representative bead, *cyan asterisks* a representative CLiP). Spd-2 beads also organise MTs that are long in interphase and short in mitosis, but they do not detectably generate CLiPs. N≥5 embryos; n≥50 beads analysed per condition. Scale bar = 10 µm.

**Figure S9.**
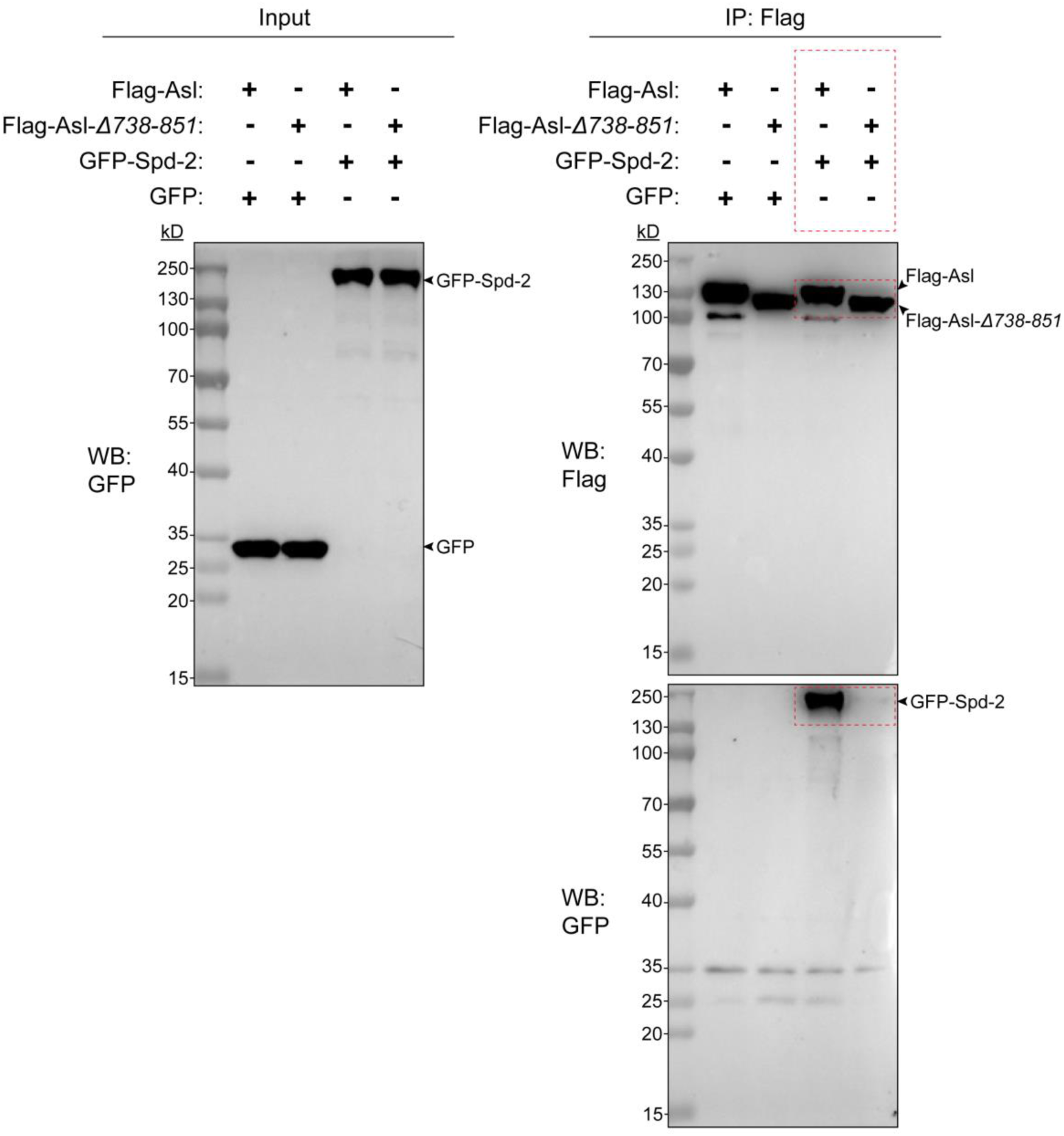
The Asl_738-851_ region is required for the interaction between Asl and Spd-2 in S2 cells. Western blots show a co-immunoprecipitation experiment from insect cells co-expressing GFP alone or GFP-Spd-2 together with either Flag-Asl-WT or Flag-Asl-Δ*738-851* (as indicated). Input lysates (left) were probed with anti-GFP to confirm expression of GFP and GFP-Spd-2. Anti-Flag immunoprecipitates (right) were probed with anti-Flag (top) to confirm equivalent recovery of Flag-Asl-WT and Flag-Asl-Δ*738-851*, and with anti-GFP (bottom) to detect co-precipitated GFP-Spd-2. GFP-Spd-2 is efficiently co-immunoprecipitated with Flag-Asl-WT but not with Flag-Asl-Δ*738-851* (*dashed red* boxes), demonstrating that residues 738-851 of Asl are required for its interaction with Spd-2. Molecular weight markers (kD) are indicated. Blots are representative examples from 2 independent experiments.

**Figure S10.**
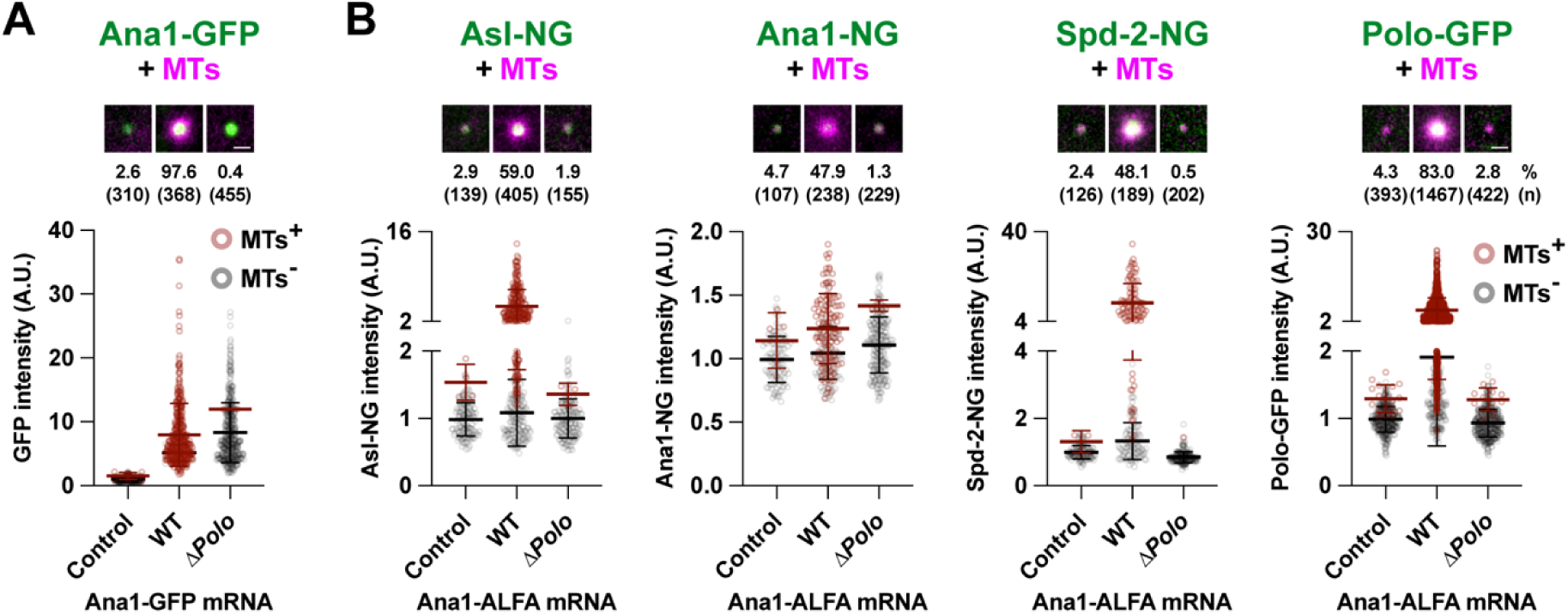
The ability of Ana1 to recruit Polo is required for Ana1-ALFA beads to recruit downstream centrosome proteins and organise MTs. (**A,B**) Images illustrate and graphs quantify the recruitment of Ana1-WT-GFP or Ana1-Δ*Polo*-GFP (*green*) to GFP-Nanobody beads (A), or of Asl-NG, Ana1-NG, Spd-2-NG or Polo-GFP (*green*) to Ana1-WT-ALFA or Ana1-Δ*Polo*-ALFA bound to ALFA-Nanobody beads (B) (as indicated). Embryos expressed Jup-mCh (MTs) (*magenta*). Each dot on the graph represents a bead, scored as either MT +ve (*brown* dots) or -ve (*grey* dots). The percentage of beads organising MTs (%) and the number of beads analysed (n) is indicated; N=7-20 embryos per condition. Ana1-Δ*Polo* is recruited to the beads at similar levels to Ana1-WT (A), but it fails to efficiently recruit Spd-2 or Polo, and fails to organise MTs (B). Perhaps unexpectedly, Ana1-Δ*Polo*-ALFA beads also recruited much less Asl and consequently failed to generate CLiPs. This confirms that Ana1 likely generates CLiPs by recruiting Asl and also suggests that Ana1’s ability to recruit Asl requires that Ana1 be able to recruit Polo. Scale bars = 2µm.

**Figure S11.**
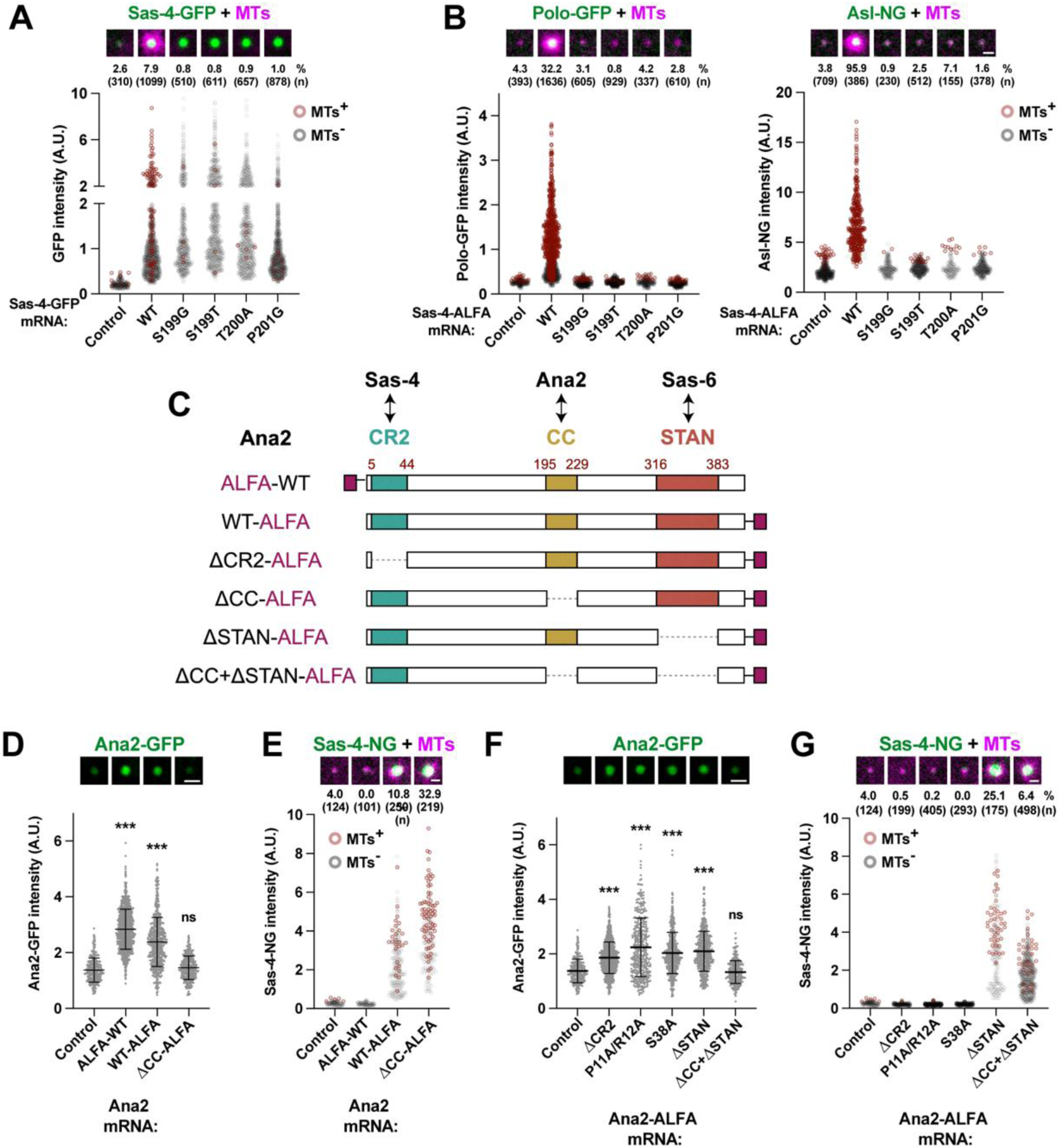
Sas-4, Ana2 and Sas-6 beads appear to engage the canonical PCM- and centriole-assembly pathways to organise MTs and generate CLiPs. (**A,B**) Images illustrate and graphs quantify the recruitment of Sas-4-WT-GFP or various point-mutant forms of Sas-4-GFP to GFP-Nanobody beads (A), or of Polo-GFP (B) or Asl-NG (C) (g*reen*) to anti-ALFA-Nanobody beads in embryos injected with mRNA encoding these proteins (as indicated). Embryos also expressed Jup-mCh (MTs) (*magenta*). N=8-27 embryos per condition. The Sas-4-GFP mutants either cannot recruit Polo (S199G and S199T) or cannot be phosphorylated on T200 by Cdk (T200A and P201G) ^3^. They were recruited to beads as efficiently as WT (A), but none recruited Polo (B) or Asl (C), and none organised MTs. (**C**) Schematic of Ana2 showing the CR2 (Sas-4-binding) ^4,5^, CC (self-association) ^6^ and STAN (Sas-6-binding) ^7,8^ domains, together with the WT and mutant Ana2-ALFA constructs tested in (D-G). The ALFA-tag was placed either N-terminally (ALFA-WT) or C-terminally (WT-ALFA and all mutants). Numbers indicate domain boundaries (aa). (**D,E**) Ana2-GFP recruitment (D) and Sas-4-NG recruitment and MT organisation (E) on anti-ALFA-Nanobody beads in embryos injected with the indicated ALFA-tagged Ana2 mRNAs. Both WT Ana2 constructs recruited Ana2-GFP, but the N-terminally tagged ALFA-Ana2 failed to recruit Sas-4 and did not organise MTs—suggesting that the N-terminal ALFA-tag prevents the adjacent CR2 domain from interacting with Sas-4, thus inhibiting MT organisation and CLiP generation. Ana2-ΔCC-ALFA beads cannot recruit Ana2-GFP, but can still recruit Sas-4-NG and organise MTs, indicating that Ana2 self-association is dispensable for these functions. (**F,G**) Assessment of Ana2-GFP recruitment (F) and Sas-4-NG recruitment and MT organisation (G) for a broad panel of Ana2-ALFA mutants that perturb Ana2’s interaction with Sas-4 or Sas-6. All the Sas-4 interaction mutants (ΔCR2, P11A/R12A, S38A) recruited Ana2-GFP but failed to recruit Sas-4 or organise MTs, whereas the mutant that cannot interact with Sas-6 (ΔSTAN) could still recruit Sas-4 and organise MTs (as could a ΔCC+ΔSTAN mutant that cannot interact with Ana2 or Sas-6). N=5-18 embryos per condition. Statistical comparisons in (D,F) against control by one-way ANOVA: *** p<0.05, ns = not significant. Scale bars = 2 µm.

**Figure S12.**
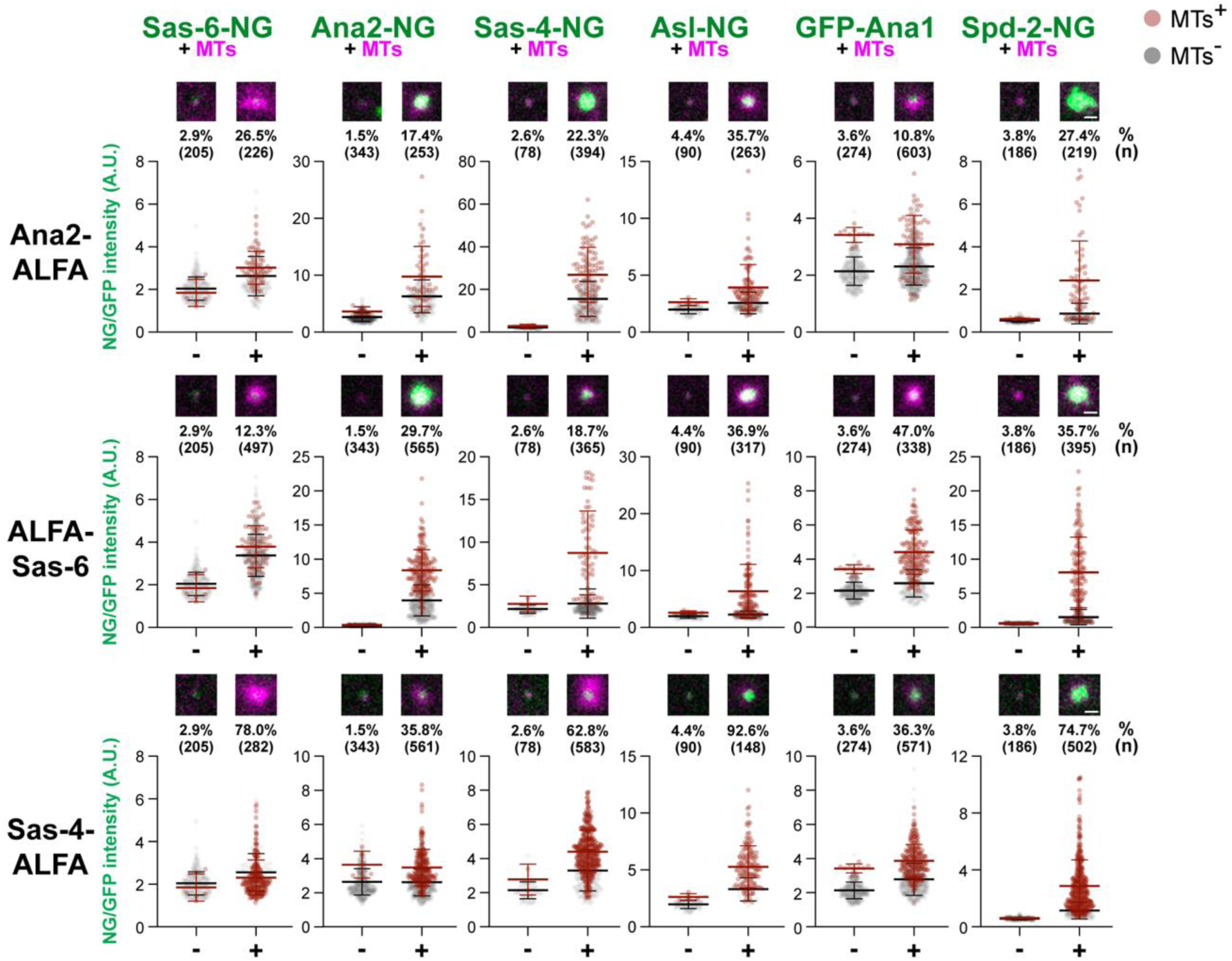
Analysis of the ability of individual core centriole and centrosome-assembly proteins to recruit other core proteins to beads to generate MTs and CLiPs. Images and graphs compare the ability of beads coupled to ALFA-tagged fusions of the cartwheel components(Ana2, Sas-6 and Sas-4, as indicated at left of each panel) to recruit various green-fluorescent fusion proteins (*green*) and generate MTs (Jup-mCh, *magenta*). Control beads are coupled to the anti-ALFA-Nanobody and injected without any mRNA. Each dot on the graph represents a bead, scored as MT +ve (*brown dots*) or -ve (*grey dots*). The percentage of beads organising MTs (%) and the number of beads analysed (n) is indicated; N=5-15 embryos per condition. This analysis broadly supports our conclusion that these cartwheel components engage the canonical assembly pathways for bead-dependent CLiP and PCM assembly (as outlined in Figure S13B). For example, Ana2 recruits relatively little Sas-6, but most of the other assembly factors, while Sas-6 recruits all of the other assembly factors. In contrast, Sas-4 recruits relatively little Sas-6 or Ana2, and more of all the other assembly factors. Importantly, all the core centriole assembly proteins should be recruited to beads at the site where CLiPs are assembling (as CLiPs contain all these proteins). This may explain why even beads that seem to recruit very little of a core protein (e.g. Sas-4-beads seem to recruit very little Sas-6 or Ana2) usually recruit at least slightly more of these proteins than controls—as some Sas-6 and Ana2 will be incorporating into the CLiPs being assembled on the bead surface. Scale bars = 2 µm.

**Figure S13.**
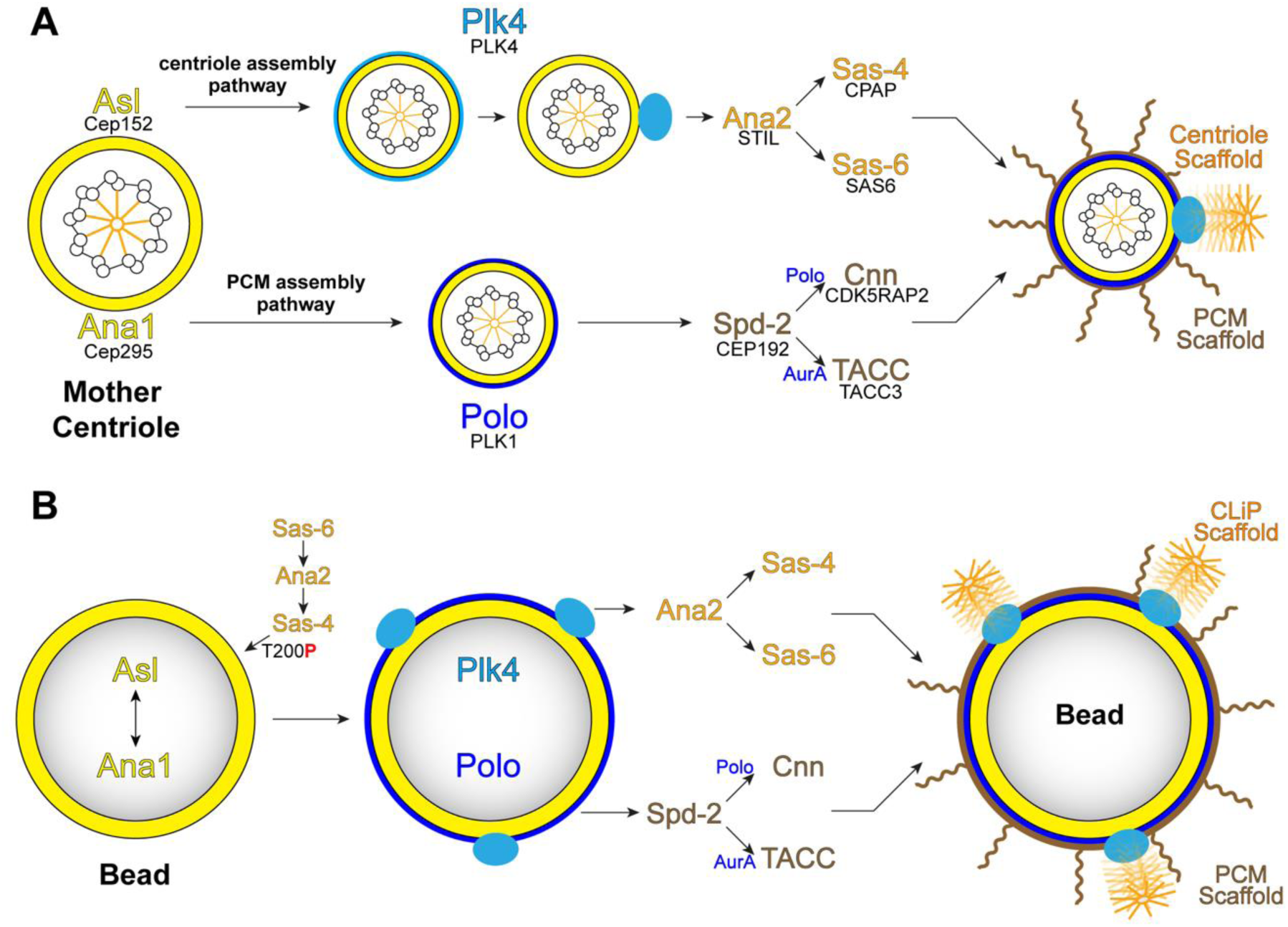
Model for how beads coupled to individual centriole assembly proteins programme CLiP and PCM assembly through the canonical pathways. Schematics compare the canonical centriole and PCM assembly pathways at a mother centriole in *Drosophila* embryos (**A**) (as shown in Figure 1B), with the proposed pathways used to generate CLiPs and PCM at the surface of synthetic beads (**B**). At mother centrioles, Asl normally recruits Plk4 to initiate daughter centriole assembly, while Ana1 normally recruits Polo to initiate PCM assembly. When coupled to beads, Asl and Ana1 can recruit each other, and so both Plk4 and Polo, to the bead surface. Plk4 can symmetry-break on the Asl-surface to generate multiple independent foci (*light blue* ovals), each of which can seed a CLiP through the canonical centriole assembly pathway. Polo is recruited to the Ana1 surface, from where it drives PCM/MT organisation in parallel via the canonical PCM assembly pathway. Beads coupled to Sas-6, Ana2 or Sas-4 feed into the same pathway via Sas-4 T200 phosphorylation— recapitulating the “daughter-to-mother conversion” step that normally occurs on the newborn daughter centriole—allowing these beads to recruit Asl and Ana1 to initiate the canonical assembly pathways.

**Figure S14.**
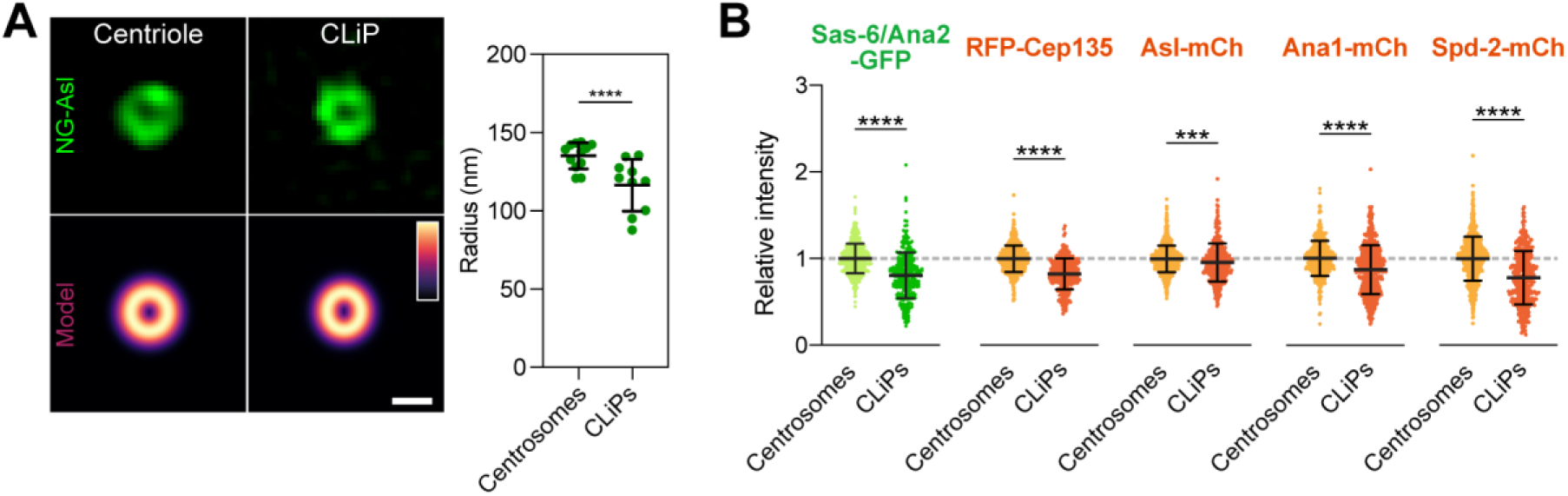
CLiPs recruit slightly less protein than centrioles and have a slightly smaller Asl-torus radius. (**A**) Super-resolution images (top) of the NG-Asl torus at a representative centriole and CLiP, together with the corresponding ring-Gaussian model fits (bottom; intensity-coded LUT shown). The graph quantifies the fitted torus radius, showing that CLiPs have a slightly, but significantly, smaller Asl-torus radius than centrioles. n=12 centrioles and n=10 CLiPs analysed. (**B**) Graphs quantify the relative fluorescence intensity of the centriole/centrosome proteins Sas-6/Ana2-GFP, RFP-Cep135, Asl-mCh, Ana1-mCh or Spd-2-mCh at centrosomes and CLiPs (normalised to the mean centrosome intensity, dashed *grey* line). CLiPs recruit slightly, but significantly, less of all these proteins than centrosomes, although there is substantial overlap between the populations. N=6-8 embryos; n > 400 centrosomes and n > 300 CLiPs per condition. Statistical comparisons by Students t-test: *** = p<0.001; **** = p<0.001. Scale bar = 200 nm. These differences are probably explained by our observation that many CLiPs are not fully mature at birth: they are associated with less centrosomal proteins and cannot duplicate (as shown in Figure S3). It has previously been shown that immature centrioles have a smaller diameter than fully mature centrioles ^9^.

**Figure S15.**
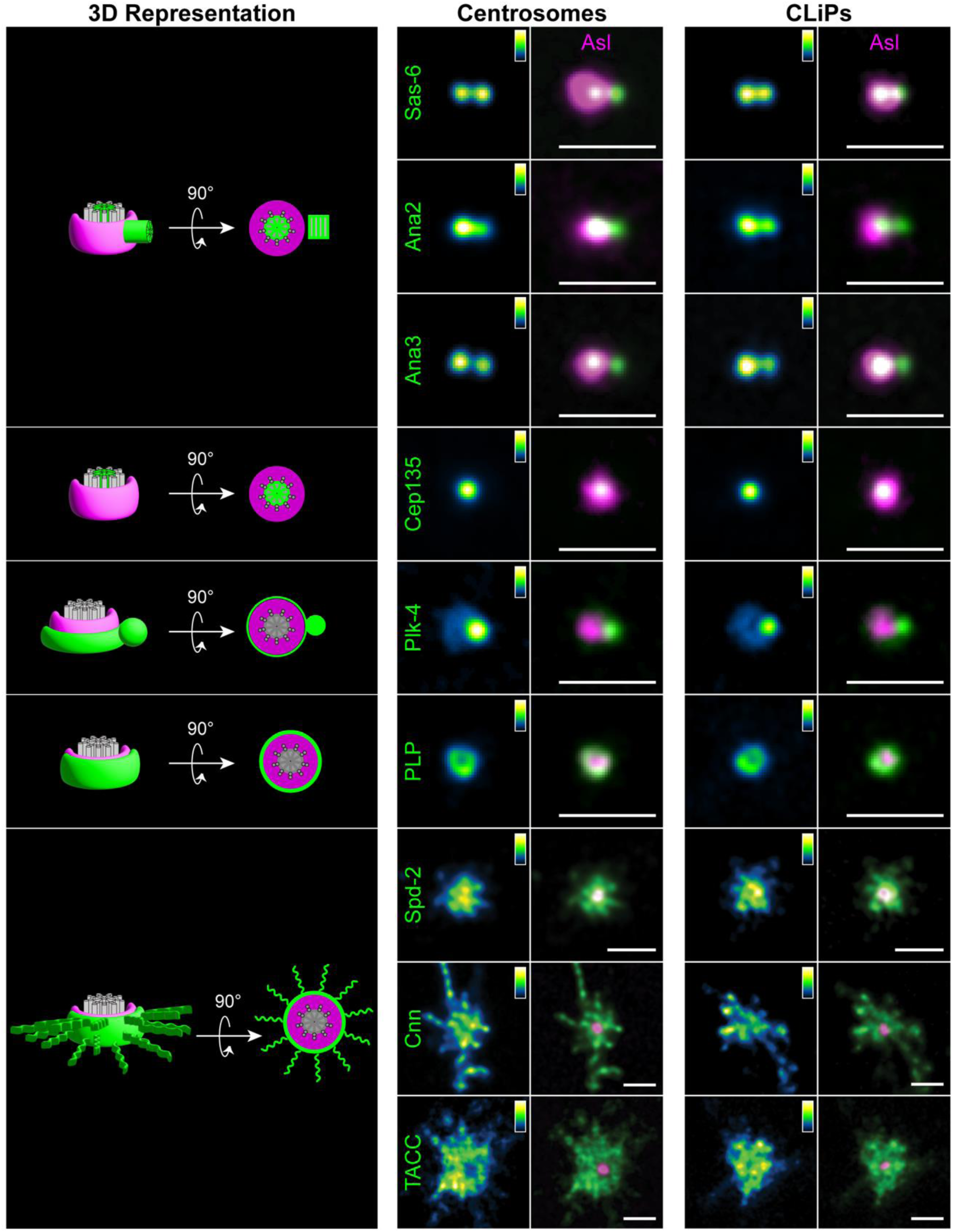
Super-resolution imaging reveals that CLiPs and centrosomes share their overall structural organisation. Schematics (left) illustrate the predicted top-down (90°-rotated) organisation of Asl (*magenta*) and each indicated centriole/PCM marker (*green*) around a mother centriole (MT barrel in *grey*) that is assembling a daughter centriole in S-phase. Super-resolution images show the top-down view of centrioles (middle) and CLiPs (right) in living embryos co-expressing Asl-mCh (*magenta*) and a NG-or GFP-fusion to each of the indicated centriole (Sas-6, Ana2, Ana3, Cep135, Plk4, PLP) or PCM (Spd-2, Cnn, TACC) proteins (*green*; displayed as a fire look-up table in single-channel panels for clarity). For each marker, CLiPs display the same spatial organisation relative to the Asl torus as *bona-fide* centrosomes, consistent with CLiPs sharing a centriole-like structural architecture. N > 3 embryos; n > 20 centrioles and n > 20 CLiPs analysed per marker. Scale bars = 500 nm.

## Video Legends

**Video S1 (related to Figure 2A)**

Beads bound to GFP-Plk4 do not generate MTs or centrosomes.

Anti-GFP-Nanobody beads were injected into an embryo expressing GFP-Plk4 (*green*) and Jup-mCh (MTs) (*magenta*). Time (mins:secs) is indicated and is relative to the first image (t=00:00), which was taken ∼20-30mins after bead injection. This video remains centred on a single bead (*black asterisk*). These beads bind high levels of GFP-Plk4 but do not generate MTs or centrosomes. Note that in these embryos GFP-Plk4 is visible on the beads but is too dim to be visualised at the centrioles. Nevertheless, this level of expression of GFP-Plk4 is sufficient to cause some centriole overduplication in the embryo ^10^, as seen by the formation of a multipolar spindle.

**Video S2 (related to Figure 2B)**

Beads bound to Polo-GFP do not generate MTs or centrosomes.

Anti-GFP-Nanobody beads were injected into an embryo expressing Polo-GFP (*green*) and Jup-mCh (MTs) (*magenta*). Time (mins:secs) is indicated and is relative to the first image (t=00:00), which was taken ∼20-30mins after bead injection. This video remains centred on a single bead (*black asterisk*). The beads bind high levels of Polo-GFP but do not generate MTs or centrosomes. In this video Polo-GFP is visible on the beads and transiently on centrosomes, kinetochores and various other mitotic structures as the embryo progresses through the cell cycle.

**Video S3 (related to Figure 3A)**

Sas-6/Ana2 beads can generate MT-organising particles when the endogenous centrosomes reach the embryo cortex.

Anti-GFP-Nanobody beads were injected into an embryo expressing Sas-6-GFP and Ana2-GFP (*green*) and Jup-mCh (MTs) (*magenta*). Time (mins:secs) is indicated and is relative to the first image (t=00:00), which was taken ∼10-20mins after bead injection. Some of the beads already organise MTs, and the embryo proceeds through two NCs (seen as pulses in the movement of the beads) before the endogenous nuclei and centrosomes start to arrive at the cortex (∼t=16:00). At this point the beads start to organise more MTs, and many of the beads that organise MTs start to generate small particles that organise MTs and move away from the bead surface (∼t=17:30). Some of the beads are initially clumped together, and many of these start to separate as the beads nucleate more MTs (∼t=23:00-26:00).

Many of the particles generated by the beads contain Sas-6/Ana2-GFP and duplicate in phase with the endogenous centrosomes (this is documented more clearly in Videos S4 and S5). After mitosis of NC13 (∼t=1:07:00), the endogenous duplicated centrosomes and many of the CLiPs separate (∼t=1:16:00), but they do not duplicate again while the embryo is cellularising during NC14. The beads and CLiPs (which are not associated with any nuclei) are gradually pushed inside the embryo by the expanding network of cellularising nuclei. This internalisation is a normal developmental programme that removes any cortical centrosomes that have lost their connection to nuclei ^1^.

**Video S4 (related to Figure 3B)**

Sas-6/Ana2 beads generate particles that duplicate in synchrony with the endogenous centrosomes.

Anti-GFP-Nanobody beads were injected into an embryo expressing Sas-6-GFP, Ana2-GFP (*green*) and Jup-mCh (MTs) (*magenta*). Time (mins:secs) is indicated and is relative to the first image (t=00:00), which was taken ∼20-30mins after bead injection. A Sas-6/Ana2 bead is highlighted (*black asterisk*) that generates several CLiPs at the end of the first round of mitosis (one of which is highlighted (*cyan asterisk*); all of these CLiPs organise MTs and duplicate in synchrony with the endogenous centrosomes after the next round of mitosis. A CLiP generated by the bead below the highlighted bead is also marked (*cyan arrowhead*). This CLiP is less bright than the others and it does not duplicate. An endogenous centrosome is highlighted for comparison (*white arrowheads*). Note that after the second round of mitosis the Sas-6/Ana2 bead generates more CLiPs, one of which is highlighted (*yellow asterisk*).

**Video S5 (related to Figure 3C)**

CLiPs can proceed through 4 rounds of duplication in synchrony with the endogenous centrosomes but, like centrosomes, they stop dividing when the embryo cellularises.

Anti-GFP-Nanobody beads were injected into an embryo expressing Sas-6-GFP, Ana2-GFP (*green*) and Jup-mCh (MTs) (*magenta*). Time (mins:secs) is indicated and is relative to the time when the CLiP of interest was first detected (t=00:00), which was ∼25-35mins after bead injection. This CLiP (*cyan box*, t=00:00) separates from the highlighted Sas-6/Ana2 bead (*black asterisk*) towards the end of mitosis in NC10. A single centrosome is highlighted at the same time for comparison (*white arrowhead*).

The duplicated centrosome separates at the end of mitosis NC10 (t=03:30) and the duplicated CLiP also separates (although this is only apparent at t=05:00). The CLiP and the centrosome proceed to duplicate and separate at the end of NC11 (t=12:30), and again at the end of NC12 (t=25:30), and again at the end of NC13 (t=42:00); so this CLiP duplicates 4 times in total. Note that after each duplication/separation event the *cyan box* and *white arrowhead* only follow one duplicated centrosome/CLiP for clarity of presentation. The embryo starts to cellularise in NC14 and the centrosomes and CLiPs no longer duplicate. Note that it becomes difficult to follow individual CLiPs as they get internalised into the yolk during the cellularisation process (see also Video S3), but the CLiP highlighted here, and the surrounding CLiPs, clearly do not duplicate during the ∼50mins they can be followed in NC14.

**Video S6 (related to Figure S3)**

CLiPs are born with varying levels of Sas-6/Ana2 fluorescence and can have different fates.

Anti-GFP-Nanobody beads were injected into an embryo expressing Sas-6-GFP, Ana2-GFP (*green*) and Jup-mCh (MTs) (*magenta*). Time (mins:secs) is indicated and is relative to the first image (t=00:00), which was taken ∼20-30mins after bead injection. A Sas-6/Ana2 bead is highlighted (*black asterisk*) that generates several CLiPs at the end of the of mitosis; an endogenous centrosome is highlighted for comparison (*white arrowheads*). The CLiPs of interest are highlighted in the corresponding Figure S3 but are not highlighted in the video for clarity of presentation. Nevertheless, it can be seen that most of the CLiPs generated have much lower levels of Sas-6/Ana2-GFP than the endogenous centrosomes (t=6:00). The brighter CLiPs (e.g., those labelled 2 and 4 in Figure S3) remain stable and their fluorescence usually increases allowing them to duplicate—either immediately, or in subsequent cycles. CLiPs born with lower levels of Sas-6/Ana2 (e.g., those labelled 1 and 3 in Figure S3) often eventually dissipate or can remain stable without duplicating. Note that even though some of the duplicating CLiPs are born with lower levels of Sas-6/Ana2 than the endogenous centrosomes, these levels often equalise in later cycles (e.g., t=24:00). We conclude that many CLiPs are born “immature” and these can either remain immature (in which case they usually ultimately dissipate) or they can gradually mature (sometimes over multiple cycles) to acquire higher levels of Sas-6/Ana2 and become duplication competent. Note also the CLiP forming the pole of a spindle (t=18:00).

**Video S7 (related to Figure 3D)**

CLiPs and beads can organise cortical actomyosin structures.

Anti-ALFA-Nanobody beads were injected with mRNA encoding ALFA-Sas-6 and Ana2-ALFA into an embryo expressing myosin light chain (MLC)-GFP (*green*) and Spd-2-mCh (centrosome marker) (*magenta*). Time (mins:secs) is indicated and is relative to the first image (t=00:00), which was taken ∼20-30mins after bead injection. The ALFA-tag is non-fluorescent, so centrosomes, beads and CLiPs are detected with the Spd-2-mCh marker. This video remains centred on a single bead (*orange asterisk*) that generates 4 CLiPs after mitosis—3 of which are highlighted, *cyan asterisks*). These CLiPs initially remain within the same MLC-domain as the bead but, after the next round of mitosis, the 3 highlighted CLiPs duplicate and organise their own domain of MLC-GFP around themselves (just as the endogenous centrosomes do—one of which, together with its progeny, are highlighted with a *white arrowhead*). Note that the cortical myosin disassembles during each round of mitosis, as actin furrows that largely lack myosin transiently form around the spindles. Also, the 4^th^ CLiP generated by the bead (not highlighted) organises a myosin cage even though it does not duplicate (t=15:00).

**Video S8 (related to Figure 3E)**

CLiPs can form spindle poles.

Anti-GFP-Nanobody beads were injected into an embryo expressing Sas-6-GFP, Ana2-GFP (*green*) and Jup-mCh (MTs) (*magenta*). Time (mins:secs) is indicated and is relative to the time when the CLiP of interest (*cyan asterisk*) could be observed to form a spindle pole (t=00:00; shown in Figure 3E), which was ∼25-35mins after bead injection. In this video, the CLiP of interest is formed at t=-15:00. It then starts to form the pole of a spindle at t=-10:30 and then does so again during the next round of mitosis (t=00:00).

**Video S9 (related to Figure 3F)**

Beads can form spindle poles.

Anti-GFP-Nanobody beads were injected into an embryo expressing Sas-6-GFP, Ana2-GFP (*green*) and Jup-mCh (MTs) (*magenta*). Time (mins:secs) is indicated and is relative to the time when the bead of interest (*black asterisk*) could be observed to form a spindle pole (t=00:00; shown in Figure 3F), which was ∼25-35mins after bead injection.

**Video S10 (related to Figure 4)**

Beads bound to Ana2-GFP can generate MTs and CLiPs.

Anti-GFP-Nanobody beads were injected into an embryo expressing Ana2-GFP from the Ubq-promotor (*green*) and Jup-mCh (MTs) (*magenta*). Time (mins:secs) is indicated and is relative to the first image (t=00:00), which was taken ∼15-30mins after bead injection. The Ana2 beads generate MTs and CLiPs.

**Video S11 (related to Figure 4)**

Beads bound to Sas-4-GFP can generate MTs and CLiPs.

Anti-GFP-Nanobody beads were injected into an embryo expressing Sas-4-GFP from the Ubq-promotor (*green*) and Jup-mCh (MTs) (*magenta*). Time (mins:secs) is indicated and is relative to the first image (t=00:00), which was taken ∼15-30mins after bead injection. The Sas-4 beads generate MTs and CLiPs.

**Video S12 (related to Figure 4)**

Beads bound to Asl-GFP can generate MTs and CLiPs.

Anti-GFP-Nanobody beads were injected into an embryo expressing Asl-GFP from the Ubq-promotor (*green*) and Jup-mCh (MTs) (*magenta*). Time (mins:secs) is indicated and is relative to the first image (t=00:00), which was taken ∼15-30mins after bead injection. For unknown reasons, the Asl-GFP beads often remain clumped together, and this video is centred on a cluster of Asl beads that generate MTs and CLiPs.

**Video S13 (related to Figure 4)**

Beads bound to GFP-Ana1 can generate MTs and CLiPs.

Anti-GFP-Nanobody beads were injected into an embryo expressing GFP-Ana1 from the Ubq-promotor (*green*) and Jup-mCh (MTs) (*magenta*). Time (mins:secs) is indicated and is relative to the first image (t=00:00), which was taken ∼15-30mins after bead injection. The Ana1 beads generate MTs and CLiPs.

**Video S14 (related to Figure 4)**

Beads bound to GFP-Sas-6 can generate MTs and CLiPs.

Anti-GFP-Nanobody beads, together with mRNA encoding GFP-Sas-6, were injected into an embryo expressing Jup-mCh (MTs) (*magenta*). Time (mins:secs) is indicated and is relative to the first image (t=00:00), which was taken ∼30-60mins after bead/mRNA injection. These Sas-6 beads generate MTs and CLiPs.

**Video S15 (related to Figure 5A)**

Sas-6/Ana2 beads require Plk4 to generate CLiPs but not to generate MTs.

Anti-GFP-Nanobody beads were injected into an embryo expressing Sas-6-GFP, Ana2-GFP (*green*) and Jup-mCh (MTs) (*red*) together with Plk4 RNAi. Time (mins:secs) is indicated and is relative to the first image (t=00:00), which was taken ∼30-45mins after bead injection. The centrosomes in this embryo (two highlighted with *white arrowheads*) had already stopped duplicating by the time the video was started, so the spindles that form around the chromatin during mitosis have either only one or no endogenous centrosomes associated with them. The embryo continues to cycle despite the lack of centrosome duplication, and the remaining centrosomes and beads continue to organise MTs that cycle in synchrony with each other. In these embryos, however, the centrioles cannot duplicate and the beads cannot generate CLiPs. Note that, due to the failure in centriole duplication, the actomyosin network that normally stops nuclei and spindles from colliding with one another is no longer organised properly. As a result, the beads in Plk4 RNAi embryo can more easily access the mitotic chromatin to form spindle poles. In this video, two beads (*black asterisks*) that are initially close together separate to form both poles of a spindle in the first round of mitosis. One of these beads then forms the pole of two spindles in the second round of mitosis.

**Video S16 (related to Figure 5D)**

Beads bound to Asl-ΔPlk4 generate MTs but not CLiPs.

Anti-ALFA-Nanobody beads, together with mRNA encoding either Asl-WT-ALFA (left video) or Asl-ΔPlk4-ALFA (right video) (that cannot recruit Plk4), were injected into embryos expressing Sas-6-NG (centriole marker) (*green*) and Jup-mCh (MTs) (*magenta*). Time (mins:secs) is indicated and is relative to the first image (t=00:00), which was taken ∼30-60mins after bead/mRNA injection. Asl beads tend to remain clumped together, and these videos show clumps of WT Asl beads organising MTs and generating CLiPs, while the clumps of Asl-ΔPlk4 beads organise MTs but do not generate CLiPs.

**Video S17 (related to Figure 5E)**

Beads bound to Asl-ΔPlk4 do not generate CLiPs.

Anti-ALFA-Nanobody beads, together with mRNA encoding either Asl-WT-ALFA (left video) or Asl-ΔPlk4-ALFA (right video) (that cannot recruit Plk4), were injected into embryos expressing Plk4-NG (*green*) and Asl-mCh (*magenta*). Time (mins:secs) is indicated and is relative to the first image (t=00:00), which was taken ∼30-60mins after bead/mRNA injection. Asl beads tend to remain clumped together, and the left video shows a cluster of WT Asl beads recruiting several small dots of Plk4-NG to their surface. These dots appear to be the source of the CLiPs that can be seen separating from the bead surface. These CLiPs initially do not recruit Plk4-NG, but become Plk4-NG positive a few minutes later, presumably as they mature and become competent to duplicate. Note that levels of Plk4-NG on the beads and on the CLiPs rise and fall in synchrony with the embryonic cell cycle, as is the case at endogenous centrioles ^11^. In contrast, Asl-ΔPlk4 beads (right video) do not organise any Plk4-NG dots and do not generate CLiPs (all the green dots visible in this video are endogenous centrioles).

**Video S18 (related to Figure 6C)**

Beads bound to Asl-Δ738-851 generate CLiPs, but do not recruit PCM.

Anti-ALFA-Nanobody beads, together with mRNA encoding either Asl-WT-ALFA (left video) or Asl-Δ738-851-ALFA (right video) (that cannot recruit Ana1, Spd-2 or Polo), were injected into embryos expressing Sas-6-NG (centriole marker, *green*) and Spd-2-mScarlet-I3 (Spd-2-I3, PCM marker, *magenta*). Time (mins:secs) is indicated and is relative to the first image (t=00:00), which was taken ∼30-60mins after bead/mRNA injection. Asl beads tend to remain clumped together, and the left video shows a cluster of Asl-WT beads recruiting Spd-2-I3 all over their surface, and Sas-6-NG to a small number of dots that appear to be the source of the CLiPs that can be seen separating from the bead surface. In contrast, Asl-Δ738-851 beads (right video) do not recruit Spd-2-I3 all over their surface but still recruit Sas-6-NG dots that generate CLiPs.

**Video S19 (related to Figure 7A)**

NG-Asl forms a torus around the barrel of mother centrioles and CLiPs.

Anti-ALFA-Nanobody beads, together with mRNA encoding ALFA-Sas-6 and Ana2-ALFA mRNA were injected into an embryo expressing NG-Asl (*green*) and Jup-mCh (MT marker) (*magenta*). Time (mins:secs) is indicated and is relative to the first image (t=00:00), which was taken ∼30-45mins after bead injection. This video remains centred on a bead that initially generates a single CLiP (*cyan asterisk*) at the same time as an endogenous duplicated centriole (*white arrowhead*) separates at the end of mitosis. The asterisk and arrowheads follow the CLiP and centrioles through two rounds of duplication. It can be seen that NG-Asl (*green*) forms a torus around the centrioles and CLiPs that can be visualised as either a ring (in an end-on view of the centriole/CLiP barrel) or a bar (in a side-on view of the centriole/CLiP barrel) on the SoRa super-resolution spinning disc system used here. The centrioles and CLiPs move around in the embryo, flipping between end-on and side-on views of the NG-Asl torus.

**Video S20 (related to Figure 7B)**

The centriole MT marker CenSpark is recruited to CLiPs shortly before they separate from the bead surface.

Anti-ALFA-Nanobody beads, together with mRNA encoding Asl-ALFA mRNA and the fluorescent centriole MT marker CenSpark (*magenta*) were injected into embryos expressing either Sas-4-NG (left video), SG-Asl (middle video) or Plk4-NG (right video) (*green*). Time (mins:secs) is indicated and is relative to the first image (t=00:00), which was taken ∼30-45mins after bead injection. These videos illustrate the incorporation timing of CenSpark relative to these centriole markers. Plk4-NG and Sas-4-NG essentially continuously mark the site of CLiP generation, and it can be seen that CenSpark only starts to accumulate at these sites shortly before the CLiPs separate from the bead surface. In contrast, SG-Asl only normally starts to incorporate into newly formed centrioles as they disengage from their mothers to become new mothers at the end of mitosis. Hence SG-Asl is not always present on the bead surface and only starts to accumulate at CLiPs shortly before they separate from the bead surface. Careful inspection reveals that CenSpark and Asl start to accumulate at CLiPs at approximately the same time, suggesting that the centriole MTs only start to form on CLiPs shortly before they separate from the bead surface.

## Materials and Methods

### Drosophila melanogaster stocks and husbandry

All *Drosophila* melanogaster stocks used in this study are listed in Table S2. Flies were maintained at 18°C or 25°C in plastic vials or bottles containing standard cornmeal-based medium composed of 0.68% agar, 2.5% yeast, 6.25% cornmeal, 3.75% molasses, 0.42% propionic acid, 0.7% ethanol, and 0.14% tegosept. For embryo collection, adult flies were transferred to cages containing fruit-juice agar plates (40% cranberry-raspberry juice, 2% sucrose, and 1.8% agar) supplemented with a drop of yeast paste. Fly work was performed using procedures designed to minimize plastic waste and improve environmental sustainability 1.

### Drosophila S2 cells

*Drosophila* S2 cells, kindly provided by Prof. Xinhua Lin (Fudan University), were cultured at 25°C in Schneider’s medium supplemented with 10% heat-inactivated fetal bovine serum, without CO2.

### Generation of transgenic and knock-in *Drosophila* lines

To generate an endogenous C-terminal Ana3-mNG knock-in line, pCFD3-Ana3-gRNA and the Ana3-mNG homology-directed-repair donor plasmid were injected at final concentrations of 0.1 µg/µl and 0.5 µg/µl, respectively, into embryos expressing Cas9 under the control of the nanos promoter (Bloomington Drosophila Stock Center stock 54591), as described previously (Port et al., 2014; Port et al., 2015). Embryo injections were performed by the University of Cambridge Department of Genetics Fly Facility. Surviving founder flies were crossed to SM5 second-chromosome balancer flies, and progeny were screened by PCR using primers located within mNeonGreen (mNG) together with primers located outside the donor homology arms. Candidate knock-in alleles were validated by PCR amplification and Sanger sequencing of both integration junctions and the modified ana3 locus. The resulting Ana3-mNG line, designated A3C6_1, was homozygous viable and fertile. All gRNA and screening-primer sequences are listed in Table S4.

The StayGold-Asl and Sas-4-StayGold transgenic lines were generated using the pNoP-eAsl-StayGold and pNoP-eSas-4-StayGold constructs, respectively. These constructs contain extended genomic regions of asl or sas-4 fused in frame to C-terminal StayGold. The constructs were introduced into the Drosophila germline using standard P-element mediated transgenesis, with embryo injections performed by the Fly Facility, University of Cambridge. Transgenic progeny were identified using the mini w+ marker, balanced as appropriate, and stable lines were established.

### Molecular cloning

#### General construction of pRNA expression plasmids

cDNA fragments encoding Sas-6, Ana2, Sas-4, Asl, and Plk4 were amplified from Drosophila cDNA, and pRNA plasmids carrying GFP, ALFA, or mNeonGreen tags were amplified as vector backbones using Phusion Flash High-Fidelity PCR Master Mix (F548, Thermo Fisher Scientific). Following amplification, DpnI (R0176, NEB) was added to each reaction and samples were incubated at 37°C for 1 h. PCR products were purified using the QIAquick PCR Purification Kit (28106, QIAGEN), assembled using NEBuilder HiFi DNA Assembly Master Mix (E5520, NEB), and incubated at 50°C for 30 min. Domain-deletion and point-mutant constructs were generated using the same PCR-based assembly strategy.

Constructs used for mechanistic analyses included an Asl mutant lacking the predicted Plk4-binding region involving Asl residues 18–54 (AslΔPlk4), Ana1ΔPolo (previously termed Ana1-ALL 2), and Sas-4 point mutants S199A, S199T, T200A, and P201G. For the Ana2 analyses shown in Figure S10, Ana2ΔCR2 lacked residues Ana25–44 and was used to disrupt the Ana2–Sas-4 interaction, Ana2ΔCC lacked residues Ana2195– 229 and was used to disrupt Ana2 dimerization, and Ana2ΔSTAN lacked residues Ana2316–383 and was used to disrupt the Ana2–Sas-6 interaction. Ana2 point mutants S38A and P11A/R12A were also used in these analyses. All ALFA-tagged Ana2 constructs used in these experiments carried the ALFA tag at the C terminus. Primer sequences used to generate these constructs are listed in Table S4, and all plasmids used in this study are listed in Table S3.

#### S2-cell expression constructs

For S2-cell experiments, pAc5.1-Spd-2-EGFP and pAc5.1-FLAG-Asl were used. The pAc5.1-FLAG-AslΔ738-851 construct was generated by PCR-based deletion followed by NEBuilder HiFi DNA Assembly using the corresponding primers listed in Table S4. Destination vectors for transgenic constructs The no-promoter C-terminal mNG Gateway destination vector pNoP-mNG-CT-DEST-attB was generated from pNoP-NeonGreen-CT-DEST by replacing portions of the 5′ and 3′ P-element sequences with an attB sequence. The parental plasmid was amplified using Q5 Hot Start High-Fidelity Master Mix (M0494S, NEB) and primers containing attB-compatible overlaps. Following DpnI digestion and agarose-gel purification, amplified fragments were assembled using NEBuilder HiFi DNA Assembly. The assembly product was transformed into ccdB-resistant E. coli, and kanamycin-resistant colonies were screened by sequencing.

The no-promoter C-terminal StayGold Gateway destination vector pNoP-StayGold-CT-DEST was generated independently from the same pNoP-NeonGreen-CT-DEST parental plasmid. The backbone was amplified using VeriFi Hot Start DNA Polymerase (PB10.43-01, PCR Biosystems) with primers oZNc-129_StayGoldF and oZNc-128_StayGoldR, which contained overlaps with a synthetic StayGold fragment. Following DpnI digestion, the PCR product was resolved by agarose-gel electrophoresis and purified using the Monarch DNA Gel Extraction Kit (T1130S, NEB). The purified backbone and synthetic StayGold fragment were assembled using NEBuilder HiFi DNA Assembly at 50°C for 1 h. The resulting plasmid was transformed into ccdB Survival competent E. coli, selected on kanamycin-containing plates, and verified by sequencing.

#### Gateway cloning of fluorescently tagged centrosomal proteins

Gateway LR recombination was used to generate Asl-StayGold, Sas-4-StayGold, Cep135-mNG, and Spd-2-mScarlet-I3 constructs. Standard reactions contained 0.75 µl pDONR entry plasmid, 0.75 µl destination plasmid, 1 µl LR Clonase II, and 2.5 µl TE in a final volume of 5 µl and were incubated at 25°C for 2.5 h. Reactions were terminated by addition of 0.5 µl Proteinase K followed by incubation at 37°C for 10 min. Reaction products were transformed into competent E. coli and selected on kanamycin-containing plates. Individual colonies were expanded, plasmid DNA was purified, and final constructs were verified by sequencing.

For pNoP-eAsl-StayGold, pDONR-Zeo-eAsl-WT-genomic-NoStop was recombined with pNoP-StayGold-CT-DEST. pNoP-eSas-4-StayGold was generated by recombining pDONR-Zeo-eSas-4-WT-genomic-NoStop with pNoP-StayGold-CT-DEST. These constructs contain extended genomic regions of asl or sas-4, respectively, fused in frame to C-terminal StayGold.

For pUbq-Cep135-mNG, pDONR-Cep135-WT-NoStop was recombined with pUbq-mNG-CT-DEST. For pUbq-Spd-2-mScarlet-I3-attB, pDONR-Spd-2-WT was recombined with pUbq-mScarlet-I3-CT-DEST. The entry and destination plasmids were each used at 0.75 µl at 100 ng/µl. The resulting pUbq-Spd-2-WT-CT-mScarlet-I3-attB construct was verified by sequencing.

To generate pNoP-eAna1(cDNA)-mNG-attB, an extended ana1 genomic fragment spanning approximately 2 kb upstream of the start codon to the final coding codon, excluding the stop codon, was first introduced into pNoP-mNG-CT-DEST-attB by Gateway LR recombination to generate pNoP-eAna1(+introns)-mNG-attB. The genomic ana1 coding region in this intermediate was then replaced with ana1 cDNA. The plasmid backbone was amplified from pNoP-eAna1(+introns)-mNG-attB using primers oZNc104 and oZNc105, and ana1 cDNA was amplified from pRNA-Ana1(WT)-mKate2 2 using primers oZNc102 and oZNc103. The two fragments were assembled using NEBuilder HiFi DNA Assembly to generate pNoP-eAna1(cDNA)-mNG-attB.

#### Ana3 CRISPR constructs

For generation of the Ana3-mNG CRISPR knock-in, complementary oligonucleotides encoding a single-guide RNA targeting the 3′ end of the ana3 coding region were phosphorylated, annealed, and ligated into BbsI-digested pCFD3 to generate pCFD3-Ana3-gRNA. The corresponding homology-directed-repair plasmid, pUC57-Ana3-mNG-HDR, contained genomic sequences flanking the 3′ end of ana3 and an in-frame C-terminal linker followed by mNG. Silent substitutions were introduced into the gRNA-recognition sequence to prevent Cas9-mediated re-cleavage of the repaired locus. The donor was synthesized in a pUC57 ampicillin-resistant backbone. Both plasmids were propagated in E. coli, purified, and verified by sequencing before embryo injection.

#### Plk4 RNAi template

For Plk4 RNA interference, primer pairs were designed in MacVector to target a 600-nt region of Plk4 cDNA and were checked for potential off-target matches using NCBI BLAST. The target region was amplified from Plk4 cDNA using Phusion Flash High-Fidelity PCR Master Mix. Following purification with the QIAquick PCR Purification Kit, a T7 RNA polymerase promoter sequence was added using a universal primer, and the resulting PCR product was purified again as described above.

### *In vitro* transcription

Plasmids used for mRNA preparation were digested and linearized with AscI (R0558, NEB) and purified using the QIAquick PCR Purification Kit (28106, QIAGEN). Linearized DNA was transcribed using the mMESSAGE mMACHINE T3 Transcription Kit (AM1348, Thermo Fisher Scientific). The resulting mRNA was purified using the RNeasy MinElute Cleanup Kit (74204, QIAGEN) and stored at −70°C. Plasmids used in this study are listed in Table S3.

For Plk4 double-stranded RNA (dsRNA) preparation, the T7-containing PCR product was transcribed using the MEGAscript T7 Transcription Kit (AM1334, Invitrogen). Transcripts were precipitated by addition of 0.1 volume of 3 M sodium acetate and 2.5 volumes of ethanol and incubated on ice for 5 min. Following centrifugation at 4°C for 10 min at full speed, the pellet was washed twice with cold 70% ethanol and dissolved in RNase-free water. Complementary single-stranded RNAs were annealed at 67.5°C for 30 min followed by slow cooling to room temperature over 60 min. The resulting dsRNA was purified as described above and stored at −70°C.

### Synthetic bead preparation

1-µm Dynabeads MyOne Streptavidin C1 (65001, Invitrogen) were washed three times in PBS containing 0.01% Triton X-100 and 0.1% BSA (PBSTB). For nanobody coupling, 100 µg of beads were incubated with 5.5 µg of biotinylated anti-GFP or anti-ALFA nanobody in a total volume of 125 µl PBSTB for at least 30 min at room temperature. Nanobody expression, biotinylation, and purification were performed as described previously 3. Beads were subsequently washed three times in PBSTB and stored at 3 mg/ml in PBSTB at 4°C for up to one week.

### Embryo collection, preparation, and microinjection

Embryos were collected from fruit-juice agar plates after a 30-min egg-laying period at 25°C. Embryos were manually dechorionated and mounted on a strip of glue in 35-mm glass-bottom dishes with 14-mm microwells (P35G-1.5-14-C, MatTek). Mounted embryos were desiccated for 5 min at 25°C and covered with Voltalef oil (H10S PCTFE, Arkema). Nanobody-coated beads were mixed with mRNA to a final bead concentration of 1 mg/ml; for control injections without mRNA, an equivalent volume of water was added instead because the mRNA stocks were prepared in water. Final concentrations in the injection mixture were 150 ng/µl for mRNA and 100 ng/µl for Plk4 dsRNA. For the Plk4-depletion experiments shown in Figure 5A, Plk4 dsRNA was co-injected with the nanobody-coated beads into embryos before nuclear cycle 10, with an equivalent volume of water used for vehicle-control injections. Embryos were incubated for a further 30 min at 25°C after injection and before imaging. During this incubation, dishes were placed on a magnet to pull the injected beads toward the embryo cortex and facilitate imaging.

### CenSpark labeling of centriole microtubules

To label centriole microtubule doublets/triplets, CenSpark 4 (Cat. CY-SC525, Cytoskeleton Inc.) was co-injected with mRNA and ALFA-nanobody-coated beads into 30-min-aged embryos at a final concentration of 100 µM in the injection mixture. The Drosophila line used for CenSpark labeling is indicated for each individual experiment. Embryos were incubated for 30 min at 22°C before imaging.

### Spinning-disk confocal microscopy

Embryos were imaged at room temperature using an Andor Dragonfly 505 spinning-disk confocal microscope with a 40-µm pinhole, mounted on a Leica DMi8 stand and controlled with Fusion software (Oxford Instruments). Excitation was provided by 488-nm and 561-nm solid-state diode lasers. Images were acquired using an HC PL APO 63×/1.40 NA oil-immersion objective and an Andor iXon Ultra 888 EMCCD camera. Z-stacks comprised 41 optical sections acquired at 0.5-µm intervals. Time interval, exposure time, laser power, EM gain, and pixel size varied between experiments and are specified for the corresponding individual experiments.

### Super-resolution spinning-disk microscopy

Super-resolution imaging was performed on a Nikon Eclipse Ti2 inverted microscope equipped with a Yokogawa SoRa spinning-disk confocal unit and a high-sensitivity sCMOS camera. Images were acquired using a 60×/1.3 NA silicone-immersion objective with silicone oil of refractive index 1.406, yielding an effective magnification of 168× or 240×. Samples were excited with 488-nm and 561-nm laser lines, and fluorescence was collected through appropriate band-pass filters in combination with a quad-band dichroic mirror.

CenSpark-labeled embryos were imaged on an Olympus IX83 microscope equipped with a Yokogawa SoRa spinning-disk unit, a 3.2× relay lens (Olympus), and a Photometrics BSI sCMOS camera (95% quantum efficiency, 6.5-µm pixels). A 60×/1.3 NA silicone-immersion objective was used with Immersol 518 F (Carl Zeiss; refractive index 1.518), giving an effective magnification of 192×. Excitation was provided by 488-nm and 561-nm QBIS lasers (Coherent), and emission was collected through 525/50-nm and 617/75-nm band-pass filters, respectively, with a 405/488/561/640-nm quad-band dichroic mirror. At least nine z-sections were acquired per image with a 0.3-µm step size. Laser power and exposure time were adjusted according to the experiment.

### SoRa reconstruction and three-dimensional deconvolution

SoRa super-resolution image reconstruction and three-dimensional deconvolution were performed in NIS-Elements (Nikon). Images were reconstructed using SoRa mode (CSU W1-SoRa modality) and subsequently subjected to fast 3D deconvolution using the Richardson-Lucy MaxIP algorithm with medium-noise settings for eight iterations.

### Measurement of Asl torus radius

The radial localization of Asl was assessed as described previously 5 using a custom Python script available at https://github.com/RaffLab/SIMcentriole-ring-measurement. For each analysis, 10–15 Asl torus structures in total were selected from three embryos and fitted with an elliptical annular Gaussian profile to determine the major and minor axes. Structures with a major-to-minor axis ratio <1.2 were considered to have sufficiently low eccentricity for analysis. Torus radius was defined as the fitted major-axis radius.

### Image processing and bead fluorescence analysis

Image stacks were processed in Fiji (ImageJ2, version 2.3.0/1.53q). Z-stacks were converted to maximum-intensity projections and corrected for photobleaching using an exponential-fit algorithm. Background signal was subtracted using a rolling-ball radius of 50 pixels. Objects were detected and their fluorescence intensities quantified using the TrackMate plugin 6. Detection thresholds were adjusted manually for each fluorescently tagged protein to account for differences in background fluorescence and to exclude unwanted background detections. For centrosome and CLiP detection, an estimated object diameter of 1 µm was used; for bead detection, an estimated diameter of 3 µm was used.

### Western blotting

For comparison of transgene and endogenous protein levels, embryos were homogenized in SDS sample buffer containing 0.25 M Tris-HCl (pH 6.8), 8% (w/v) SDS, 20% β-mercaptoethanol, 40% (v/v) glycerol, and 0.08% bromophenol blue. Protein corresponding to eight embryos was loaded per lane on 3–8% Tris-acetate precast SDS-PAGE gels (Invitrogen, Thermo Fisher Scientific) and transferred to 0.2-µm nitrocellulose membranes (#162-0112, Bio-Rad). Membranes were blocked in 1× TBS containing 4% milk powder and 0.1% Tween-20 and probed with primary antibodies, followed by HRP-conjugated secondary antibodies.

Primary antibodies included rabbit anti-Sas-6 7, rabbit anti-Ana2 8, rabbit anti-Asl 9, rabbit anti-Sas-4 9, rabbit anti-Cnn 10, mouse anti-actin (1:500; Sigma, AB_476730) (all diluted 1:500 in blocking bugger). HRP-conjugated donkey anti-rabbit IgG (NA934V, lot 17876631, Cytiva Life Sciences) and sheep ECL anti-mouse IgG (NA931V, GE Healthcare) were used at 1:3000. Chemiluminescent signals were developed using SuperSignal ECL substrate (34580, Thermo Fisher Scientific) and recorded on film.

### S2-cell transfection and co-immunoprecipitation

For co-immunoprecipitation, S2 cells were seeded in 60-mm dishes at 2 × 106 cells per dish and transfected with 3 µg of the indicated plasmids (FLAG-Asl or FLAG-AslΔ738-851 together with Spd-2-GFP) using QIAGEN Transfection Reagent (Cat. 301427) according to the manufacturer’s instructions. Seventy-two hours after transfection, cells were harvested, washed in ice-cold PBS, and lysed for 1 h on ice in 50 mM Tris-HCl (pH 7.4), 150 mM NaCl, 0.5% NP-40, and protease inhibitors. Lysates were cleared by centrifugation at 14,000 × g for 30 min at 4°C, and an input fraction was retained.

Cleared lysates were incubated with anti-FLAG affinity beads (SA042005, Smart-Lifesciences) for 2 h at 4°C with rotation. Beads were washed three times with wash buffer consisting of lysis buffer diluted fivefold, and bound proteins were eluted in SDS sample buffer. Eluates and input samples were analyzed by western blotting using rabbit monoclonal anti-GFP (1:5000; AE078, ABclonal) and mouse monoclonal anti-FLAG (1:5000; AE005, ABclonal) primary antibodies, followed by HRP-conjugated anti-rabbit IgG (1:5000; M21002S, Abmart) or HRP-conjugated anti-mouse IgG (1:5000; M21001S, Abmart).

### Protein structure prediction and visualization

The full-length amino acid sequences of Drosophila melanogaster Plk4 (Sak-PA isoform) and Asterless (Asl-PA isoform) were obtained from FlyBase. Structural predictions were performed using AlphaFold3 through the AlphaFold Server with the full-length sequences of both proteins as input. Predictions were evaluated using server-provided confidence metrics, including the interface predicted TM-score (ipTM) and predicted aligned error (PAE). The full-length prediction yielded an ipTM of 0.32, while the predicted interface showed low PAE values.

A more focused prediction was subsequently performed using Asl residues 1–70 and the Plk4 kinase/PB1-PB2 region (residues 370–620). This prediction yielded an ipTM of 0.74 and identified a well-defined interface involving Asl residues 18–54, consistent with the Plk4-binding region tested experimentally. Predicted structures were visualized and analyzed using UCSF ChimeraX version 1.7.1, which was also used to generate structural figures.

### Operational definition and scoring of CLiPs

For the analyses in this study, a centrosome-like particle (CLiP) was defined as a discrete particle that emerged from and detached from the surface of a programmed bead and organized its own detectable microtubule array. CLiPs were followed through successive nuclear cycles to determine whether they subsequently duplicated.

Scoring of microtubule-organizing activity Microtubule-organizing activity was quantified using the fluorescence-analysis workflow described above under “Image processing and bead fluorescence analysis.” For each experiment, the relevant control population was specified for that comparison, and an object was classified as microtubule-positive when its microtubule-marker signal exceeded the control mean by more than two standard deviations.

### Spd-2 and Plk4 pulse quantification

Spd-2 and Plk4 fluorescence intensities were quantified using the workflow described above under “Image processing and bead fluorescence analysis.” Fluorescence traces were aligned to the embryonic cell cycle using Jupiter-mCherry as the microtubule marker, which allowed entry into M phase to be identified directly from the characteristic changes in microtubule organization.

### Statistical analysis

All statistical analyses were performed in GraphPad Prism, with details of the statistical tests provided in the corresponding figure legends.

---

**Table S1.** Reagents and resources used in this study.

| REAGENT or RESOURCE | SOURCE | IDENTIFIER |
| --- | --- | --- |
| **Antibodies** |  |  |
| Rabbit anti-Sas-6 | 1 | N/A |
| Rabbit anti-Ana2 | 2 | N/A |
| Rabbit anti-Asl | 3 | N/A |
| Rabbit anti-Sas-4 | 3 | N/A |
| Rabbit anti-Cnn | 4 | N/A |
| Mouse anti-actin | Sigma | RRID:AB_476730 |
| Rabbit monoclonal anti-GFP | ABclonal | Cat#AE078 |
| Mouse monoclonal anti-FLAG | ABclonal | Cat#AE005 |
| HRP-conjugated donkey anti-rabbit IgG | Cytiva Life Sciences | Cat#NA934V; lot 17876631 |
| Sheep ECL anti-mouse IgG | GE Healthcare | Cat#NA931V |
| HRP-conjugated anti-rabbit IgG | Abmart | Cat#M21002S |
| HRP-conjugated anti-mouse IgG | Abmart | Cat#M21001S |
| **Chemicals, Peptides, and Recombinant Proteins** |  |  |
| Dynabeads MyOne Streptavidin C1 | Invitrogen | Cat#65001 |
| Biotinylated anti-GFP nanobody | 5 | N/A |
| Biotinylated anti-ALFA nanobody | 5 | N/A |
| CenSpark | Cytoskeleton Inc. | Cat#CY-SC525 |
| Voltalef grade H10S oil | Arkema | H10S PCTFE |
| SuperSignal ECL substrate | Thermo Fisher Scientific | Cat#34580 |
| Anti-FLAG Affinity Beads | Smart-Lifesciences | Cat#SA042005 |
| Phusion Flash High-Fidelity PCR Master Mix | Thermo Fisher Scientific | Cat#F548 |
| DpnI | New England Biolabs | Cat#R0176 |
| QIAquick PCR Purification Kit | QIAGEN | Cat#28106 |
| NEBuilder HiFi DNA Assembly Master Mix | New England Biolabs | Cat#E5520 |
| Q5 Hot Start High-Fidelity Master Mix | New England Biolabs | Cat#M0494S |
| VeriFi Hot Start DNA Polymerase | PCR Biosystems | Cat#PB10.43-01 |
| Monarch DNA Gel Extraction Kit | New England Biolabs | Cat#T1130S |
| mMESSAGE mMACHINE T3 Transcription Kit | Thermo Fisher Scientific | Cat#AM1348 |
| RNeasy MinElute Cleanup Kit | QIAGEN | Cat#74204 |
| MEGAscript T7 Transcription Kit | Invitrogen | Cat#AM1334 |
| Ascl | New England Biolabs | Cat#R0558 |
| QIAGEN Transfection Reagent | QIAGEN | Cat#301427 |
| MatTek 35-mm glass-bottom dish, 14-mm microwell | MatTek | Cat#P35G-1.5-14-C |
| 3–8% Tris-acetate precast SDS-PAGE gels | Invitrogen / Thermo Fisher Scientific | Cat#EA0375BOX |
| 0.2-μm nitrocellulose membrane | Bio-Rad | Cat#162-0112 |
| **Experimental Models: Cell Lines** |  |  |
| Drosophila S2 cells | Gift from Prof. Xinhua Lin, Fudan University | N/A |
| **Software and Algorithms** |  |  |
| Fiji (ImageJ2), version 2.3.0/1.53q | NIH | https://imagej.net/ |
| TrackMate | Tinevez et al., 2016 / ImageJ | https://imagej.net/plugins/trackmate/ |
| GraphPad Prism | GraphPad | Version 10 |
| AlphaFold Server / AlphaFold3 | Google DeepMind / Isomorphic Labs | Accessed 13 July 2026; seed 1850772731 |
| UCSF ChimeraX | UCSF | Version 1.7.1 |
| Custom Asl ring-measurement script | Raff Lab | https://github.com/RaffLab/SIMcentriole-ring-measurement |

**Table S2.** Drosophila fly lines used in this study.

| FLY LINE | SOURCE | IDENTIFIER |
| --- | --- | --- |
| D. melanogaster: Jupiter-mCherry | 6 | N/A |
| D. melanogaster: Ubq-GFP-Plk4 | 1 | N/A |
| D. melanogaster: ePolo-GFP | 7 | N/A |
| D. melanogaster: eAna2-GFP; eSas6-GFP | This study | N/A |
| D. melanogaster: Ubq-Spd-2-mCherry | 8 | N/A |
| D. melanogaster: eSas-4-mCherry | 9 | N/A |
| D. melanogaster: Ubq-Asl-mCherry | 10 | FlyBase ID:<br>FBal0343645 |
| D. melanogaster: Ubq-RFP-Cnn | 11 | N/A |
| D. melanogaster: Ubq-mCherry-TACC | 5 | N/A |
| D. melanogaster: GFP-Sqh (Myosin regulatory light chain) | 12 | N/A |
| D. melanogaster: eAna2-GFP | 13 | N/A |
| D. melanogaster: Ubq-Ana2-GFP | This study | N/A |
| D. melanogaster: eSas-4-GFP | 3 | N/A |
| D. melanogaster: Ubq-Sas-4-GFP | 1 | N/A |
| D. melanogaster: eAsl-GFP | 14 | FlyBase ID:<br>FBtp0040947 |
| D. melanogaster: Ubq-Asl-GFP | 2 | N/A |
| D. melanogaster: eAna1-GFP | 14 | N/A |
| D. melanogaster: Ubq-GFP-Ana1 | 15 | N/A |
| D. melanogaster: eSas-6-GFP | 13 | FlyBase ID:<br>FBtp0131375 |
| D. melanogaster: Ubq-GFP-Sas-6 | 1 | N/A |
| D. melanogaster: eAna1-NG; Jupiter-mCherry | This study | N/A |
| D. melanogaster: Spd-2-NG | 16 | N/A |
| D. melanogaster: eSas-6-NG, Jupiter-mCherry | 17 | N/A |
| D. melanogaster: eAsl-mCh; Plk4-NG | This study | N/A |
| D. melanogaster: eAsl-NG | 7 | N/A |
| D. melanogaster: eSas-6-NG | This study | N/A |
| D. melanogaster: Spd-2-mScarletl3 | This study | N/A |
| D. melanogaster: NG-eAsl | 7 | N/A |
| D. melanogaster: StayGold-Asl | This study | N/A |
| D. melanogaster: Asl-mCherry; Ubq-GFP-Cep97 | This study | N/A |
| D. melanogaster: Sas-4-StayGold | This study | N/A |
| D. melanogaster: Ubq-Spd-2-GFP | 18 | N/A |
| D. melanogaster: Sas-4-NG; Jupiter-mCherry | This study | N/A |
| D. melanogaster: Ana2-NG; Jupiter-mCherry | This study | N/A |
| D. melanogaster: Ubq-RFP-Cep135 | 19 | N/A |
| D. melanogaster: Ubq-Ana1-mCherry | 15 | N/A |
| D. melanogaster: eAna2-NG | 17 | N/A |
| D. melanogaster: Ana3-NG | This study | N/A |
| D. melanogaster: Cep135-NG | This study | N/A |
| D. melanogaster: Asl-mCherry; CP110-GFP | This study | N/A |
| D. melanogaster: PLP-GFP | 20 | N/A |
| D. melanogaster: NG-Cnn | 16 | N/A |
| D. melanogaster: NG-TACC | 5 | N/A |
| D. melanogaster: ePlk4-NG | 21 | N/A |

**Table S3.** Plasmids used in this study.

| PLASMID | SOURCE | IDENTIFIER |
| --- | --- | --- |
| pAc5.1-Spd2-EGFP | This study | N/A |
| pAc5.1-FLAG-Asl | This study | N/A |
| pAc5.1-FLAG-AslΔ738–851 | This study | N/A |
| pNoP-mNG-CT-DEST | This study | N/A |
| pNoP-mNG-CT-DEST-attB | This study | N/A |
| pNoP-StayGold-CT-DEST | This study | N/A |
| pUbq-mNG-CT-DEST | This study | N/A |
| pUbq-mScarlet-I3-CT-DEST-attB | This study | N/A |
| pDONR-Zeo-eAsl-WT-genomic-NoStop | This study | N/A |
| pDONR-Zeo-eSas4-WT-genomic-NoStop | This study | N/A |
| pDONR-Cep135-WT-NoStop | This study | N/A |
| pDONR-Spd2-WT | This study | N/A |
| pNoP-eAsl-StayGold | This study | N/A |
| pNoP-eSas4-StayGold | This study | N/A |
| pUbq-Cep135-mNG | This study | N/A |
| pUbq-Spd2-mScarlet-I3-attB | This study | N/A |
| pNoP-eAna1(+introns)-mNG-attB | This study | N/A |
| pRNA-Ana1(WT)-mKate2 | <sup>15</sup> | N/A |
| pNoP-eAna1(cDNA)-mNG-attB | This study | N/A |
| pCFD3-Ana3-gRNA | This study | N/A |
| pUC57-Ana3-mNG-HDR | This study | N/A |

**Table S4.**
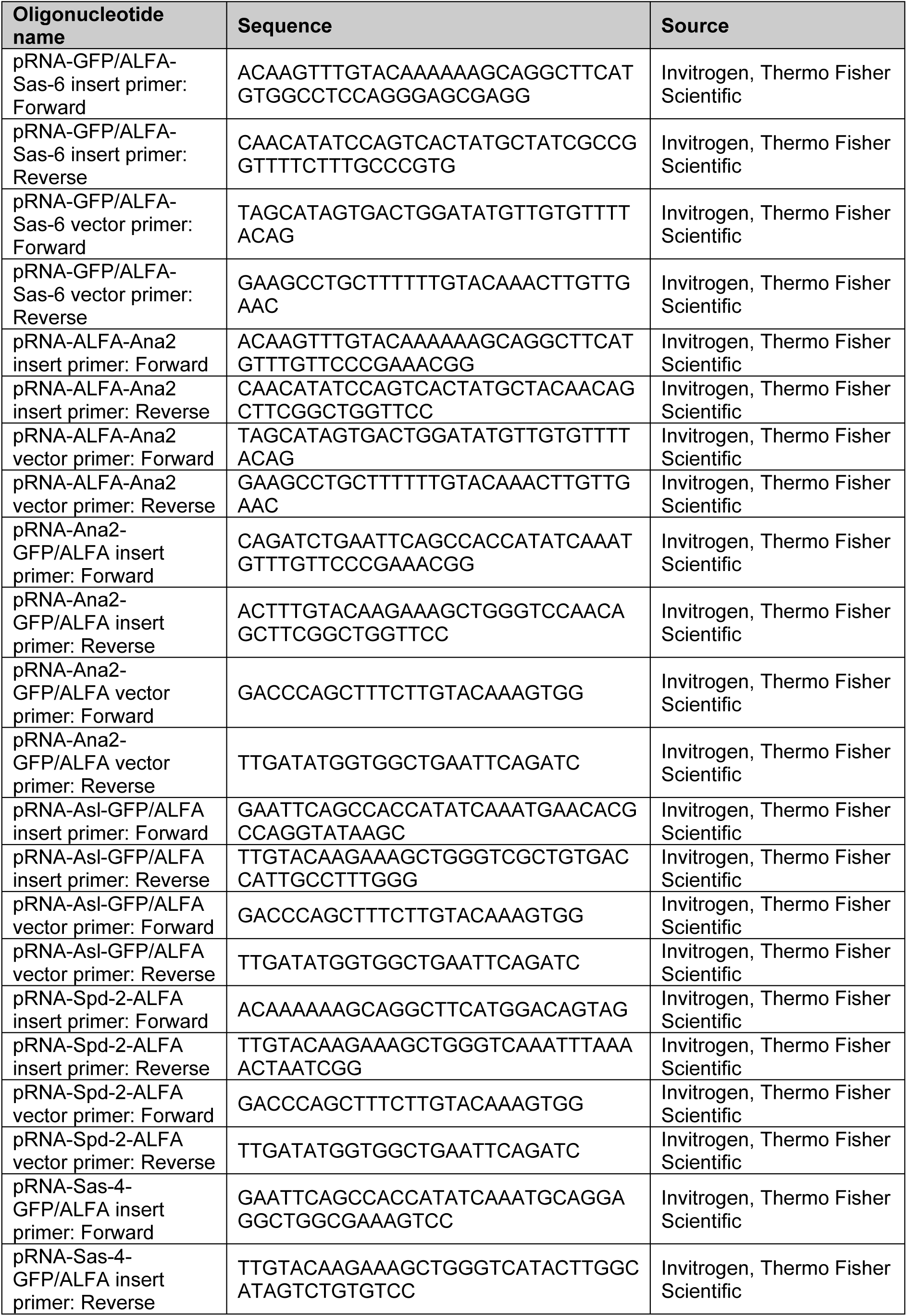
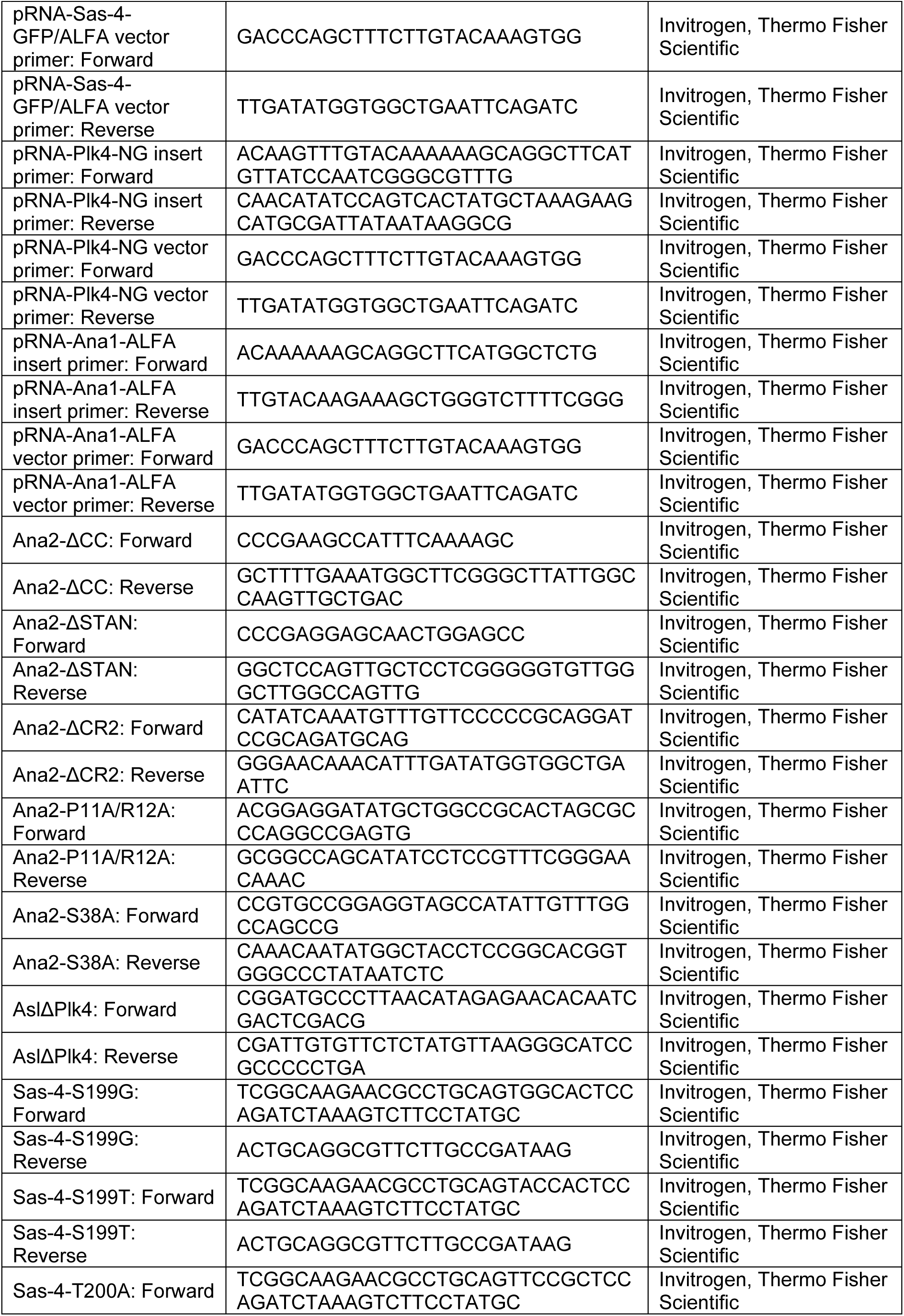
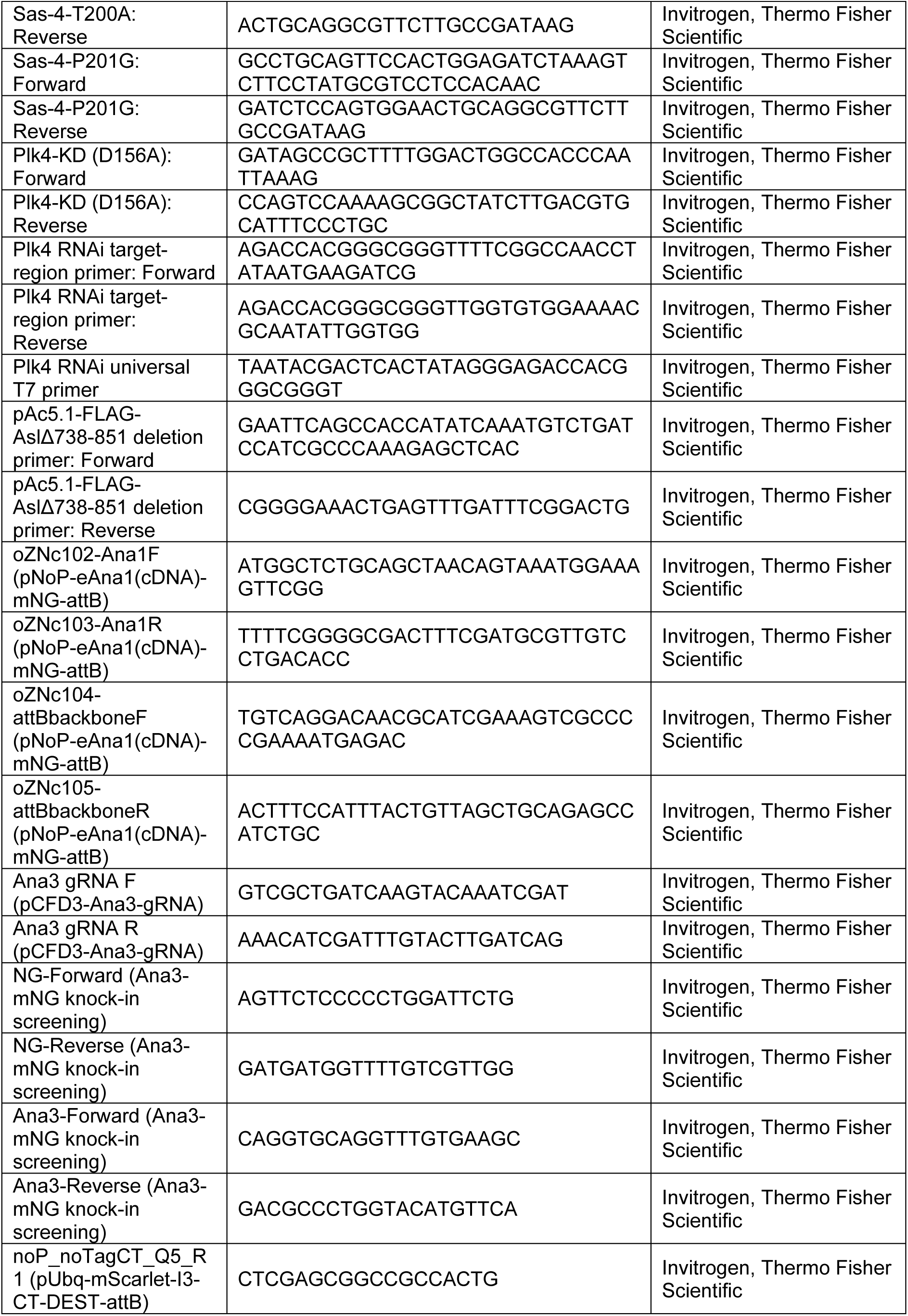
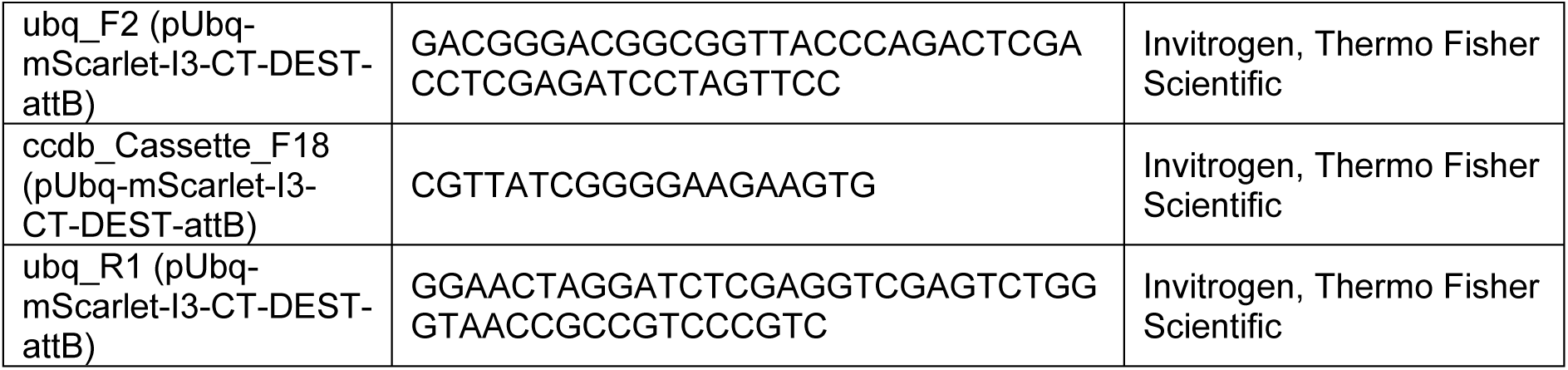
Oligonucleotides used in this study.

